# How Fisherian runaway halts: Genomic architecture determines the long-term evolutionary consequences of sexual selection

**DOI:** 10.64898/2026.11.04.764172

**Authors:** Kotaro Kagawa, Masakado Kawata

## Abstract

Sexual selection explains the evolution of exaggerated sexual displays, but less obvious is why biased mate preferences—a requirement for sexual selection—evolve. Fisher (1930) proposed that biased mate preferences can spark self-reinforcing evolutionary process (runaway) that drives their own evolution. This occurs because mate choice imposes sexual selection on sexual displays and links them genetically to mate preferences, producing indirect selection on the preferences themselves. While mathematical models have demonstrated the plausibility of the runaway theory, they rely on several idealized assumptions. Consequently, it remains poorly understood how runaway halts once populations deviate from these idealized conditions as a consequence of evolution via runaway, and how the genetic architecture underlying sexual displays influences the resulting evolutionary dynamics. Here, we address these gaps using two complementary approaches. First, we review and synthesize existing theories to classify the mechanisms that halt runaway into four categories, based on which idealized assumptions of Lande’s (1981) classic model are violated during evolution via runaway. Second, we conduct individual-based evolutionary simulations varying the genetic architecture of sexual displays from conventional additive to omnigenic inheritance models. Our results show that genetic architecture determines how runaway process halts, thereby governing long-term evolutionary dynamics under sexual selection.

## Introduction

Most theoretical models in evolutionary biology either do not explicitly consider the genetic basis of traits (e.g., game-theoretic models) or adopt simplified assumptions about it (e.g., infinitesimal models of quantitative genetics and most population-genetic models featuring a few genetic loci). A common rationale justifying such simplifications is that when evolution is driven by selection, the ultimate phenotypic outcome of evolution is expected to be determined mostly by selective pressures, rather than being strongly dependent on the underlying genetics. On the other hand, recent genomic studies are increasingly revealing more complex genetic underpinnings of diverse traits (Barghi *et al*. 2020; Fagny & Austerlitz 2021; Signor & Nuzhdin 2018). For example, there is a growing recognition that many complex traits are “omnigenic” (Box 1), influenced directly by a limited set of genes involved in the developmental process of the traits (hereafter, “core genes”) and indirectly by numerous regulatory regions affecting the expression patterns of the core genes (hereafter, “peripheral genes”) (Boyle *et al*. 2017; Liu *et al*. 2019). Despite the advances in our understanding of the genetic architecture of traits, whether and how the predictions from models with simplified genetic assumptions are robust under more nuanced genetic models has rarely been studied.

In the present study, we focus on Fisher’s runaway theory as a case in which evolutionary dynamics can qualitatively change depending on genetic architecture of traits. Fisher’s runaway theory was developed to explain the evolution of sexually selected ornaments—such as nuptial coloration, courtship songs, and courtship dances—that offer no apparent survival benefit. The evolution of such costly sexual displays can be explained by intersexual selection, a type of sexual selection driven by mate choice by the opposite sex (to simplify discussion, hereafter we consider the case where males serve as the courting sex and females serve as the choosing sex). Less obvious, however, is why animals often show biased mate preferences that drive intersexual selection. To date, several mechanisms have been proposed to explain how biased mate preferences can evolve by providing females with direct or indirect fitness benefits (reviewed in Kokko *et al*. 2006; Kuijper *et al*. 2012; Rosenthal 2017). Fisherian selection is one of the mechanisms conferring indirect fitness benefits of mate choice. As argued by Fisher (1930), female mate choice creates both sexual selection on male displays and a genetic correlation (linkage disequilibrium) between alleles encoding male displays and female mate preferences. For example, offspring of females that prefer ornamented males tend to inherit both alleles encoding the male ornament (inherited from father) and alleles encoding the preference for ornamented males (inherited from mother). Because of the genetic correlation, sexual selection on male displays also causes indirect selection on female mate preferences (i.e., Fisherian selection). From a female’s perspective, Fisherian selection is equivalent to an increase in the average mating success of sons conferred by alleles for attractive sexual displays inherited from the selected males, which is often referred to as the “sexy son” benefit of mate choice. Importantly, the evolution of a stronger mate preference via Fisherian selection in turn reinforces sexual selection and thus the Fisherian selection itself. Hence, Fisherian selection can cause a cycle of self-reinforcing evolution of female mate preferences—Fisher (1930) termed this process “runaway”. Later, Lande (1981) and Kirkpatrick (1982) validated the runaway theory using mathematical models. While they formulated Fisher’s verbal arguments using different modeling frameworks—quantitative -and population-genetics models—they both supported the same conclusion that a runaway process can occur under certain conditions. Since then, many subsequent theoretical studies have deepened our understanding of the evolutionary dynamics driven by the runaway process.

Current theories of Fisherian runaway, however, are subject to two major limitations. First, we know little about how the genetic architecture of traits may affect evolutionary dynamics driven by runaway, even though recent genomic studies are revealing the complex genetic basis of sexually selected traits (Blankers & Shaw 2024; Chenoweth & McGuigan 2010; Merrill *et al*. 2024; Orteu & Jiggins 2020). This is largely because most existing models of runaway adopt extremely simple genetic assumptions. For example, population-genetic models of sexual selection, including Kirkpatrick’s (1982) model, typically assume a single or a few biallelic loci for each trait. In contrast, most quantitative-genetic models of sexual selection, including Lande’s (1981) model, assume infinitely many loci with infinitesimal additive effects and that genetic variance of traits and genetic covariance among them (i.e., the G-matrix) are kept constant during evolution. In reality, genetic variance of sexual display and genetic covariance between sexual display and mate preference can change during evolution, which can alter the feasibility of runaway process itself. Such situations, however, have rarely been studied.

Second, we still lack a synthetic understanding of long-term evolutionary consequences of the runaway process, even though what we observe in nature most likely represents relatively stable states in long-term dynamics. The classical models of Lande (1981) and Kirkpatrick (1982) did not intend to predict long-term evolutionary dynamics. In Lande’s (1981) model, runaway process never stops once it starts. As Lande (1981) himself mentioned, this is an unrealistic prediction resulting from simplifying assumptions of the model. On the other hand, Kirkpatrick’s (1982) model considered the fate of single mutant alleles for a male display and female mate preference; thus, predicting long-term evolution with spontaneous mutations was outside the scope of this model. Thus, although these classical models demonstrated that runaway can drive the evolution of biased mate preferences by making random mating evolutionarily unstable, they did not clarify long-term evolutionary outcomes of runaway. Subsequent models with different biological assumptions have shown that runaway process can lead to various long-term evolutionary dynamics. Importantly, some models have suggested that evolutionary dynamics can qualitatively change depending on genetic assumptions, but these findings have not yet been synthesized into a unified, systematic understanding.

Here, we address these issues by providing (1) a review of existing theories regarding long-term evolutionary dynamics caused by runaway sexual selection and (2) individual-based simulations to investigate how genetic basis of sexual displays can influence long-term evolutionary outcomes of runaway. We start by providing an overview of Lande’s (1981) model, which we consider the reference model for long-term evolutionary dynamics caused by runaway sexual selection with spontaneous mutations. Then, based on a review of theoretical studies, we argue that the mechanisms for halting the runaway process can be categorized based on which idealized assumptions of Lande’s (1981) model are violated as a consequence of evolution via runaway. This also provides a categorization of long-term evolutionary outcomes of runaway. Finally, using an individual-based model in which the genetic architecture of male sexual displays can be controlled by parameters, we explore the conditions for each category of evolutionary outcome of runaway.

### Overview of Lande’s (1981) model

Lande’s (1981) quantitative-genetic model considers a male sexual display and female mate preference as quantitative traits that are controlled by an infinite number of genetic loci with infinitesimal additive effects. The model incorporates the population-level effects of female mate choice to generate (1) sexual selection on males and (2) genetic covariance between the male display and female mate preference. The indirect selection on female mate preference (i.e., Fisherian selection) is calculated as the product of selection gradient on male display and the additive genetic covariance between male display and female mate preference. Female mate preference receives no direct selection. These assumptions yield the following formulation of evolutionary dynamics (Kokko *et al*. 2006; Kuijper *et al*. 2012; Lande 1981; Mead & Arnold 2004):

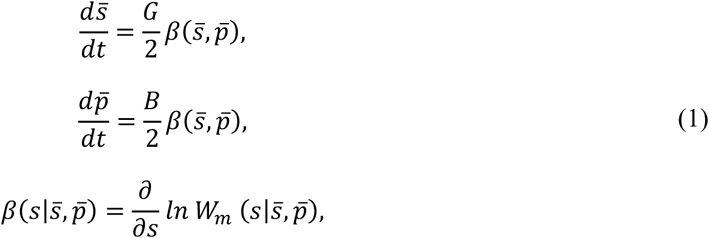

where 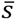 and 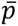 are mean values of the male display (*s*) and the female mate preference (*p*), respectively. Both *s* and *p* can take either positive or negative values. Parameters *G* and *B* are additive genetic variance of male display and additive genetic covariance between male display and female mate preference, respectively. The function *W* represents the fitness function of males, and *β* represents the selection gradient along the male display trait (*s*). Since female mate preference (*p*) is not subject to direct selection, its evolution is fully determined by the indirect selection generated via genetic covariance. Thus, the direction (sign) of evolution of 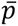 is always the same as that of 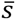.

Male fitness *W* depends on both sexual and natural selection on the male display. Sexual selection depends on how female mate preference is modeled. Under the “psychophysical preference” model in Lande (1981), 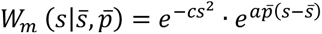. The first term expresses stabilizing natural selection, and the second term expresses sexual selectin. Note that we adopt a slightly modified parameterization of Lande’s (1981) original model. The optimal value of the stabilizing natural selection—the least costly male display—is 0, and *c* is the intensity of the stabilizing selection. The “psychophysical preference” model formulates female mate preference using an exponential function, where *p* = 0 corresponds to random mating, and positive and negative values of *p* correspond to preferences for males with positive and negative extreme values of *s*, respectively. Thus, when *p* > 0, the relative reproductive fitness of a male increases exponentially with the distance of his sexual display from the average sexual display in the population (*s* − 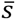). Under this model, females express open-ended mate preferences where they prefer more extreme male displays with intensity *ap*.

This model has infinitely many equilibrium points on the line 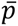 = 2*c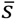*⁄*a* (Figure 1a), where sexual selection is balanced by the stabilizing natural selection (i.e., *W*_m_ (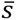, 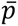) = 0). Runaway occurs if the genetic covariance (*B*) is large relative to the genetic variance of the male display (*G*); specifically, *B*/*G* > 2*c*/*a* (Figure 1c). All equilibrium points are evolutionarily unstable under this condition. In this case, sexual selection on males drives substantial evolutionary shifts in female mate preference via indirect selection. A shift in female mate preference then leads to a shift of the optimal value of male display s* at which sexual selection under the current value of 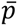 is balanced by stabilizing natural selection (i.e., *s*^∗^ *suc*ℎ *t*ℎ*at W*_m_ (*s*^∗^|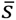, 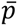) = 0). Runaway occurs because s* moves faster than average male trait 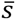. Because the evolution of the male display cannot catch up with the moving optimal value, evolution does not reach stasis during the runaway process. Graphically, the ratio of genetic covariance to variance (*B*/*G*) corresponds to the slope of evolutionary trajectories in the (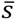, 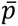) plane and 2*c*/*a* is the slope of the equilibrium line. When the slope of evolutionary trajectories is greater than that of the equilibrium line, evolutionary trajectories move away from the equilibrium line. Runaway renders random mating (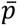 = 0) inherently evolutionarily unstable, enabling the spontaneous evolution of biased mate preferences. This holds true even if female mate preference is subject to direct stabilizing selection in favor of random mating, unlike in cases where equilibrium points are evolutionarily stable (Box 2). In Lande’s (1981) model, the runaway process never stops once initiated. This unrealistic prediction stems from the simplifying assumptions of the model. In real organisms, the runaway process may halt when evolutionary changes driven by runaway cause the conditions for runaway to be violated.

**Figure 1.**
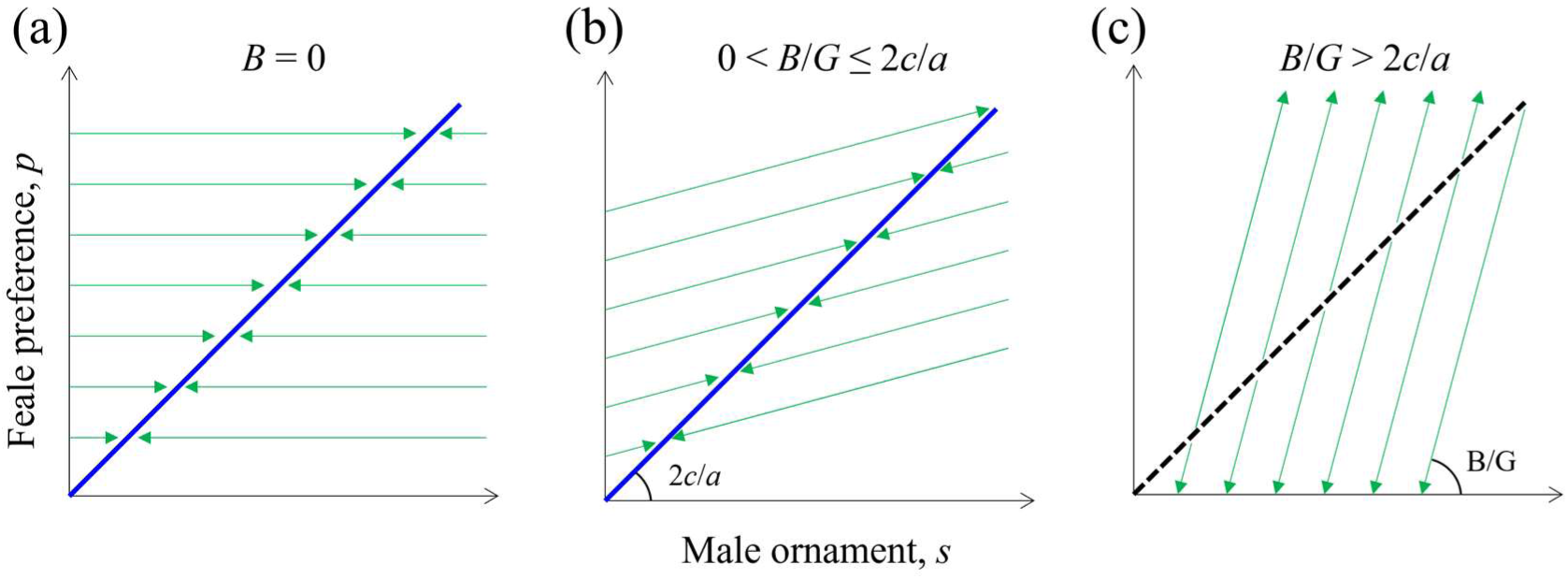
Schematic phase diagram of evolutionary dynamics in Lande’s (1981) model. The horizontal and vertical axes show population means of the male display (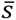) and female mate preference (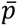), respectively. (a) Evolutionary dynamics in the absence of genetic covariance between the male display and female mate preference (*B* = 0). Since neither direct nor indirect selection affects female mate preference, the slope of the evolutionary trajectories (green lines) is zero in the (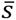, 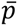) space. For each female mate preference value 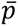, there is a stable equilibrium of the evolutionary dynamics of male display 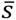, which collectively constitutes a line in the (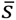, 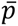) space (blue solid line). (b) Evolutionary dynamics with a weak genetic covariance. The slope of the evolutionary trajectories (green lines) is positive, but all equilibrium points are still stable. (c) Evolutionary dynamics with a sufficiently strong genetic covariance to cause runaway (*B*/*G* > 2*c*/*a*; see main text). All the equilibrium points become evolutionarily unstable (black dotted line).

#### Idealized assumptions in Lande’s (1981) model and a classification of mechanisms that halt the runaway process

In Lande’s (1981) model, the runaway process never stops once it starts because of several idealized assumptions of the model. In this section, we argue that the runaway process halts when one of the idealized assumptions in Lande’s (1981) model becomes a poor approximation of the system as a result of evolutionary changes caused by the runaway process. This also means that mechanisms by which runaway halts can be categorized based on which idealized assumption is violated.

Lande’s (1981) model makes the following idealized assumptions. (A) Infinite population: the population size is infinitely large. (B) Unlimited opportunities for mating: there is no upper limit to the number of times that individual males can mate, and females do not suffer from any cost of searching for a mate. (C) Fixed G-matrix: the additive genetic variance-covariance matrix (G-matrix) for male sexual display and female mate preference is constant during evolution. (D) Unlimited phenotypic range: there are no upper and lower limits for possible phenotypic values of traits. These assumptions simplify the model without disrupting the mechanisms of runaway, but none of them hold in nature.

Below, we review previous theoretical findings about what may occur when each of the idealized assumptions of Lande’s (1981) model becomes a poor approximation of the system as a result of runaway evolution. Our review mainly focuses on models assuming open-ended mate preferences such as the “psychophysical preference” model of Lande (1981), because most previous quantitative-genetic models in which runaway occurs under wide parameter ranges assumed open-ended mate preference functions. We do not aim to provide a comprehensive review of existing theories of runaway. Rather, our review focuses on models with relatively simple assumptions that allow direct comparison with Lande’s (1981) model; thus, we do not review population-genetic models of Fisherian runaway. Additionally, we do not discuss the effects of space, although several theoretical studies have demonstrated that long-term evolutionary dynamics can be qualitatively changed in the presence of spatial structure (Day 2000; M’Gonigle *et al*. 2012; Payne & Krakauer 1997).

### (A) Runaway can lead to evolutionary suicide in finite populations

One possible evolutionary consequence of the runaway process in finite populations is the extinction of the population (i.e., evolutionary suicide). This can occur because the evolution of increasingly costly male displays through runaway process reduces the average survival fitness of males, which increases the risk that males go extinct due to demographic stochasticity in finite populations. Even in infinite populations, runaway can lead to population extinction if evolution in a male sexual display leads to a maladaptive evolution of its homologous organ in females, thereby reducing the population growth rate (Lande & Arnold 1985).

### (B) The loss of selection gradient can halt runaway

In Lande’s (1981) model, runaway never stops once it starts, because equilibrium points constitute a straight line in the (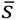, 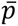) space (Figure 1). This is, however, a special case relying on specific model assumptions. Small modifications of biological assumptions from Lande’s (1981) model that slightly alter the fitness function *W* of Equation 1 mostly cause evolutionary equilibrium points to form a curve rather than a line (e.g., Iwasa & Pomiankowski 1995; Kirkpatrick *et al*. 1990). For instance, Iwasa & Pomiankowski (1995) considered a case where the cost of bearing a sexual display steeply increases when the intensity of the sexual display is large, which seems to be a plausible assumption for many organisms. Under this condition, the slope of the equilibrium curve increases with the distance from the origin (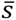 = 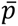 = 0). Whenever the slope of the equilibrium curve increases with |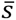| (i.e., the curve is concave up in the first quadrant), the evolutionary trajectory of runaway that moves away from the origin eventually crosses the equilibrium curve (Figure 2b). In this case, runaway process halts after reaching an equilibrium point, unlike in Lande’s (1981) model. At equilibrium, sexual selection for a costly male display and natural selection against it are balanced. Thus, the selection gradient on the male display is zero, causing Fisherian selection on females to vanish as well.

**Figure 2.**
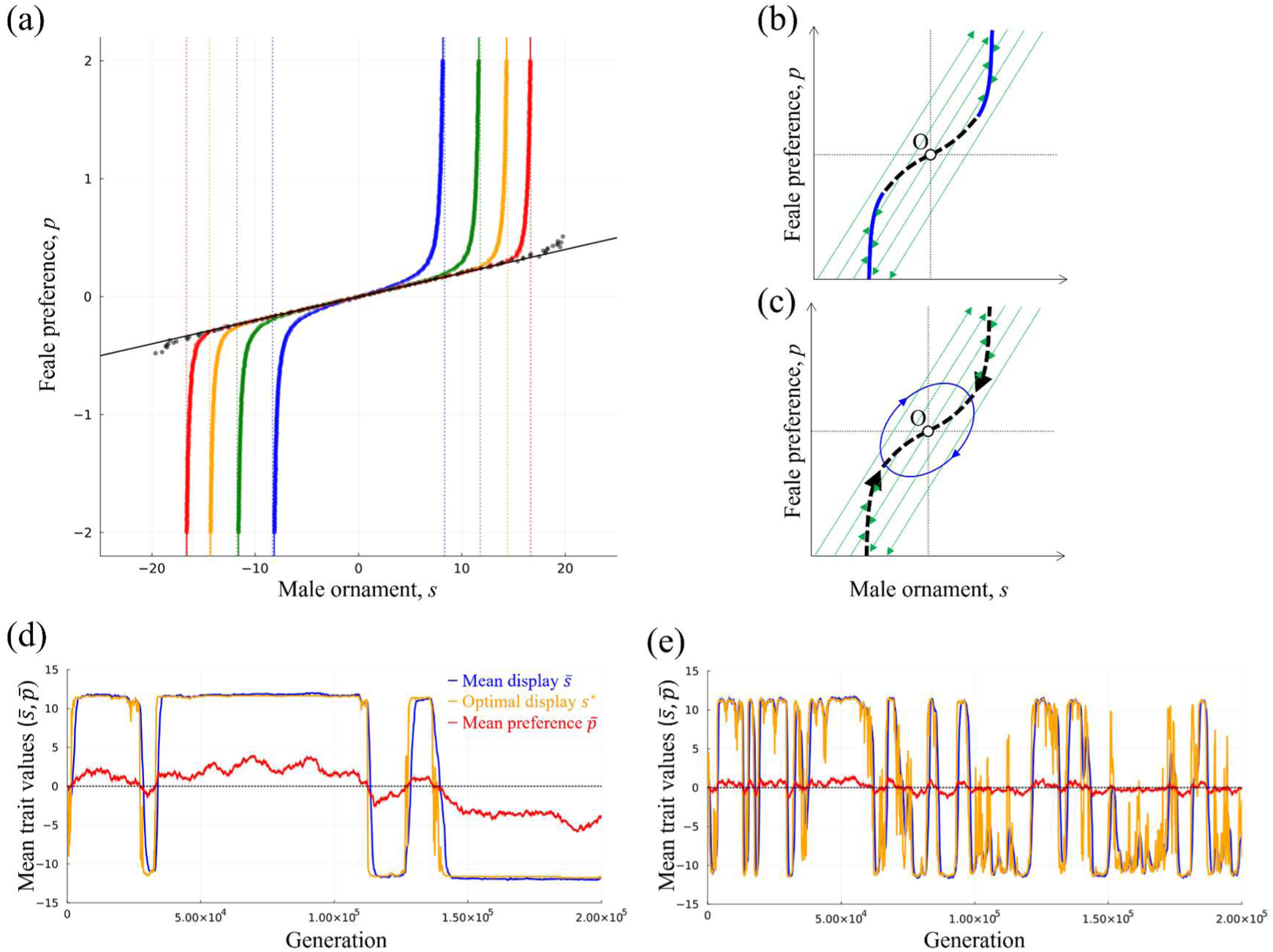
Evolutionary dynamics with finite male mating capacities. (a) Equilibrium values of the male display estimated by evolutionary simulations over 10,000 generations under fixed values of female mate preference (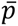). The sexual display was assumed to be controlled by 1,000 additive loci. The value of 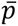 was varied from -2 to 2 in increments of 0.01. Simulations under different values of the male mating capacity (*M*) are shown in different colors (blue: *M* = 2; green: *M* = 4; yellow: *M* = 8; red: *M* = 16; black: *M* = ∞). Other parameters were set to their default values (Table 1). As expected, the slope of the equilibrium curves around the origin was the same as that of the equilibrium line in Lande’s (1981) model (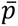 = 2*c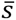*⁄*a*). Additionally, under finite male mating capacities, the equilibrium value of the male display 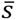 saturated at a value at which the male survival probability was 1/*M* due to the costs of bearing a sexual display (dashed lines). We fitted a cubic spline curve to the simulation results under each finite value of *M* in order to construct a formula for estimating the optimal value of the male display from the female mate preference value (solid lines). (b) Schematic phase diagram of evolutionary dynamics under a finite male mating capacity and no direct selection on female mate preference. While equilibrium points around the origin point were evolutionarily unstable (black dashed line), those distant from the origin point were neutrally stable (blue solid line). (c) Schematic phase diagram of evolutionary dynamics under a finite male mating capacity and direct stabilizing selection on female mate preference. As in panel (b), the runaway process renders the origin (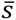 = 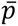 = 0) evolutionarily unstable. Simultaneously, however, direct natural selection on female mate preference destabilizes the otherwise evolutionarily stable part of the equilibrium curve. The absence of stable equilibrium points results in cyclic evolutionary dynamics in the (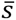, 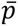) space. (d) Simulation example for the situation in panel (b). Population means of the male display (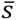; blue) and female mate preference (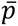; red) are shown across generations. Colored areas show the ranges from the 5th to 95th percentiles for *s* and *p*, although they are very narrow in this simulation example. The yellow line shows the dynamics of the optimal male display value under the female mate preference at each time point. Parameters: *N* = 300; *N* = 0; *N* = 0; *M* = 4; *θ* = 0.005; *K* = 1000. (e) Simulation example for the situation in panel (c). Parameters: *N* = 100; *N* = 0; *N* = 0; *M* = 4; *θ* = 0; *K* = 1000. Other parameters were set to their default values (Table 1).

**Table 1.** Parameters of the individual-based simulation model.

| **Definition** | **Symbol** | **Default** | **Alternative** |
|---|---|---|---|
| Number of core genes for sexual display | *N<sub>c</sub>* | 100 | 20, 40, 60, 80, 120,160, 200 |
| Number of trans-regulations per core gene | *N<sub>r</sub>* | 160 | 10, 20, 40, 80, 120, 160, 200 |
| Number of peripheral genes | *N<sub>p</sub>* | 2000 |  |
| Number of loci for mate preference | - | 200 |  |
| Phenotypic effect of core gene | *ε* | 0.5 |  |
| Male mating capacity | *M<sub>max</sub>* | 4 | 2, 4, 8, 16, ∞ |
| Number of males that each female encounter | *N<sub>encount</sub>* | 100 |  |
| Cost of sexual display | *c* | 0.01 |  |
| Search cost | *θ* | 0.01 |  |
| Intensity of mate choice | *a* | 1 |  |
| Carrying capacity | *K* | 5000 |  |
| Female fecundity | - | 100 |  |
| Recombination rate/bp | *r* | 10<sup>-8</sup> |  |
| Mutation rate/locus | *μ* | 10<sup>-5</sup> |  |
| Length of each locus (bp) | - | 1000 |  |
| Number of chromosomes | - | 10 |  |
| Length of each chromosome (bp) | - | 2<sup>8</sup> |  |

We note that it can generally be expected that the slope of the equilibrium curve increases with the distance from the origin (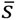 = 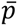 = 0) whenever males cannot mate with infinitely many females (Figure 2a shows individual-based simulations validating this argument). It is true that males have the potential to mate with many females in many organisms, and the assumption that males have unlimited chances of mating well approximates such a situation. However, whenever there is an upper limit on male reproductive fitness, the intensity of sexual selection for more extreme male displays (i.e., the marginal gain of male fitness caused by a unit increase in the intensity of sexual display) must saturate as the intensity of male display increases through the runaway process. For example, consider the case where individual males cannot mate more than 100 times. When male survival rate drops to 1/100 or less due to the cost of bearing a highly exaggerated sexual display, reproductive success of all the surviving males approaches 100. A further increase of female mate preference does not increase the intensity of sexual selection, because male fitness cannot be increased beyond this upper limit. Saturation of the intensity of sexual selection causes the slope of equilibrium curve to increase with the intensity of male display |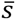| (in the first quadrant of 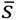-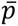 space).

What follows after runaway evolution reaches a stable equilibrium point? Two common evolutionary forces can diminish the stable persistence of exaggerated male displays, thereby leading to cyclic evolutionary dynamics. First, genetic drift in finite populations can cause slow and random evolutionary drift on the curve of stable equilibria. This can eventually bring the population out of the stable part of the equilibrium curve, which causes either runaway evolution back toward a stable equilibrium or a rapid evolution resulting in the loss of the costly male display (Figure 2d shows an individual-based simulation validating this). The latter evolutionary process is driven by an evolutionary feedback loop that is similar to the runaway process. When genetic drift causes a slight reduction of female mate preference (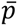) from an equilibrium point, sexual selection for the costly male display becomes slightly weaker than the natural selection against it. Because of the genetic correlation, a net selection against the costly male display generates an indirect selection against female preference for males with costly displays. This indirect selection drives the evolution of weaker female mate preference, which further weakens the sexual selection for the costly male display. During this process, females with less biased preferences gain an indirect benefit through the increase of son’s viability owing to alleles encoding less costly displays inherited from mating partners. After the loss of a costly male display, the runaway evolution occurs again, because the least costly male display (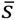 = 0) is evolutionarily unstable under the conditions where runaway process can initiate.

Second, direct fitness costs of mate preference can also hinder the stable persistence of male displays that have evolved through runaway (De Jong & Sabelis 1991; Kirkpatrick 1985; Pomiankowski *et al*. 1991). Female mate preference may receive direct stabilizing natural selection in favor of weaker preferences (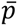 = 0; i.e., random mating) when there are costs of spending time and energy for evaluating males. Additionally, strong mate preferences can induce the risk of missing the opportunities of reproduction (i.e., the wallflower effect; De Jong & Sabelis 1991; Kokko & Mappes 2005). Introducing even a weak direct selection on the female mate preference to the model of Equation 1 removes all the equilibrium points except for the origin (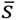 = 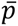 = 0). This is because while selection gradient on the male display becomes 0 only when *β*(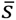, 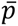) = 0, under this condition, selection gradient on the female mate preference can become 0 only when 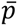 = 0 given that the indirect selection *Bβ*(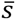, 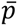)/2 = 0. Under such conditions, the runaway process making 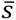 = 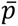 = 0 evolutionarily unstable can drive a continual cycle of evolution (Iwasa & Pomiankowski 1995), where evolution and collapse of an extreme sexual display repeat (Figure 2c, e).

### (C) The loss of genetic variance and covariance can halt runaway

Lande (1981) analyzed his model under the assumption that the additive genetic variance-covariance matrix (G-matrix) does not change during evolution. In finite populations with finite numbers of genetic loci, G-matrix changes over time due to selection, non-random mating, and genetic drift that affect allelic variation in loci and linkage disequilibrium among them. When G-matrix changes during evolution, components of the condition for the occurrence of runaway (*B*/*G* > 2*c*/*a*) change over time. Runaway can halt when this condition becomes unsatisfied during evolution.

For example, the runaway process can generate strong sexual selection that depletes male genetic variance (Nichols & Butlin 1989; Tomkins *et al*. 2004). A reduction in male display variance (*G*) in turn reduces the genetic covariance between the male display and female mate preference (*B*) by limiting females’ ability to distinguish males with different phenotypes in mate choice. In an extreme case where the genetic variance of the male display *G* drops to zero due to intense sexual selection, genetic covariance *B* also becomes zero, thereby eliminating Fisherian selection. Therefore, the runaway process driving the evolution of a strongly biased female mate preference can create conditions in which no selective pressure maintains the biased preference. This is one form of the so -called lek paradox. After the loss of the Fisherian selection, female mate preference evolves randomly due to genetic drift. Alternatively, if female mate preference is subject to a stabilizing selection, females evolve to lose the biased mate preference (Figure 3b). Accordingly, the occurrence of the lek paradox also leads to repeated evolution and collapse of costly male displays.

**Figure 3.**
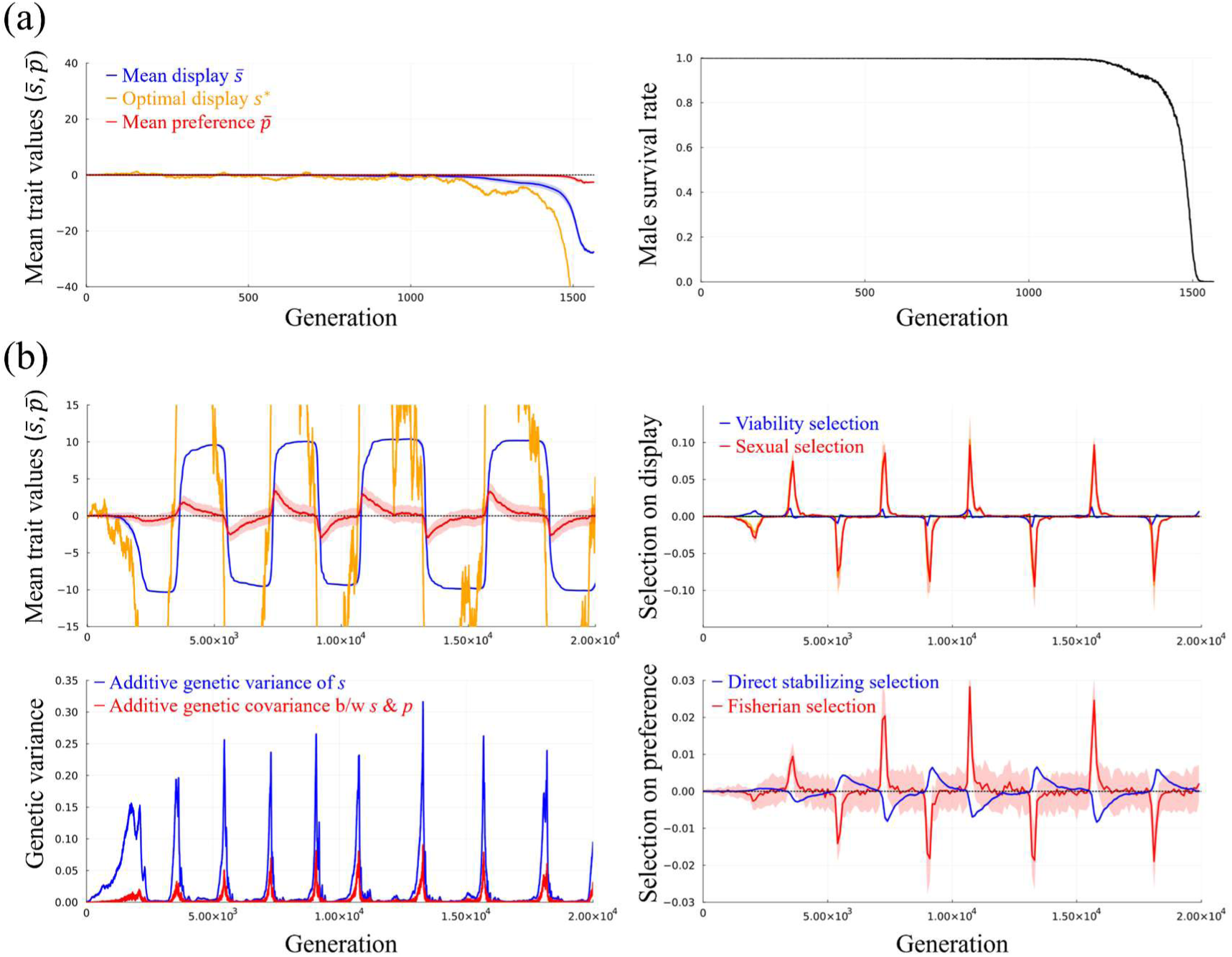
Examples of evolutionary dynamics with a male display controlled solely by loci with simple additive effects. (a) Simulation example in which runaway led to evolutionary suicide. Left: evolutionary dynamics of the population means of the male display (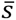; blue) and female mate preference (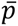; red). Colored areas show the ranges from the 5th to 95th percentiles for *s* and *p* (the ranges were very narrow in this panel). The yellow line shows the dynamics of the optimal male display value. Right: male survival rate at each generation. Parameters: *N* = 300; *N* = 0; *N* = 0; *M* = ∞; *θ* = 0.005; *K* = 10000. (b) Simulation example in which runaway led to the loss of genetic variation, causing cyclic evolutionary dynamics. Top left: evolutionary dynamics. Bottom left: the dynamics of the additive genetic variance of the male display (blue) and the additive genetic covariance between the male display and female mate preference (red). Top right: selection pressures on the male display. Blue and red lines show selection differentials on *s* in males caused by stabilizing natural selection before maturation and sexual selection after maturation, respectively. Selection differentials were grouped into 100-generation bins, with the mean values displayed for each bin. Colored areas show the ranges from the 25th to 75th percentiles of selection differentials within the bins (the ranges were very narrow in this panel). Bottom right: selection pressures on the female mate preference. The blue line shows the selection differential on *p* in females caused by direct stabilizing selection. The red line shows the selection differential on *p* in males caused indirectly by sexual selection on the male display *s* combined with the genetic correlation (i.e. Fisherian selection). Parameters: *N* = 50; *N* = 0; *N* = 0; *M* = ∞; *θ* = 0.005; *K* = 10000. Other parameters were set to their default values (Table 1).

It has been argued that the loss of genetic variation in sexually selected traits can be avoided if the mutation rate is high. More specifically, the genic capture hypothesis posits that genetic variation in sexually selected traits can be maintained when their phenotypes depend on overall condition (e.g., health or vigor), which is influenced by numerous genetic loci (Tomkins *et al*. 2004). However, virtually nothing is known about precise conditions for avoiding the loss of genetic variation during the runaway process.

### (D) Runaway can halt at stable equilibrium points with mutation bias

Genetic, developmental, and physical constraints limiting the possible range of male phenotypes can also affect the long-term evolutionary outcome of the runaway process. In Lande’s (1981) model, there are no upper and lower limits to phenotypes, allowing the runaway process to continue forever. If the ranges of possible phenotypes are limited, the runaway process cannot continue when the phenotype of male sexual display approaches a boundary. While physical constraints must always exist, genetic and developmental constraints may also exist generally at least on the timescale of microevolution where phenotypic evolution is mainly driven by slight modifications of existing genetic and developmental mechanisms. The presence of such limits to possible phenotypes is inseparable from the occurrence of mutation biases near the boundaries of the possible phenotypic range, because de novo mutations to increase (decrease) the phenotype of a trait cannot occur in genotypes with the upper (lower) limit phenotype of the trait. This fact is equivalent to the presence of a mutation bias that tends to degrade extreme phenotypes of male displays, which has been assumed in the models of Pomiankowski et al. (1991) and Iwasa & Pomiankowski (1995).

When a runaway evolution halts near a boundary of possible phenotypes, the evolutionary dynamics can reach a stable equilibrium state where sexual selection is balanced by stabilizing natural selection plus mutation bias (Pomiankowski et al. 1991). As we have seen above, runaway evolution can halt after the male display reaches an optimal value at which the selection gradient on the male display is zero. Pomiankowski et al. (1991) showed that the runaway process can halt before the selection gradient on males becomes zero when there is a mutation bias that tends to degrade extreme phenotypes of male displays. In this case, a costly male display can be stably maintained in an equilibrium state where sexual selection for more extreme phenotype is balanced by stabilizing natural selection plus the effect of mutation bias towards less extreme phenotypes. Unlike cases where sexual selection is balanced by only stabilizing natural selection, the selection gradient on males, *β*(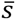, 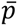), does not become zero in this equilibrium. Thus, the indirect selection for a biased female mate preference does not vanish as well.

However, the models of Pomiankowski et al. (1991) and Iwasa & Pomiankowski (1995) assumed an infinite population with a fixed G-matrix, making it unclear whether the stable maintenance of a costly male display with Fisherian selection and mutation bias is feasible under realistic conditions. In the following section, we confirm that such a maintenance can occur under certain biologically feasible conditions using individual-based simulations.

Another caveat is that the above conclusion depends on the assumption that mutations degrading a sexual display also reduce the fitness costs of bearing it. This assumption is plausible when the costs are caused by increased predation or conspecific male aggression, or when mutations tend to silence metabolic pathways allocating resources to the expression and maintenance of exaggerated sexual displays. However, this assumption does not hold when mutation bias only reduces the efficiency at which resources are converted to sizes of sexual displays without reducing resource demand for developing them, and when the primary source of the costs is resource investment to sexual displays. Dashtbali & Henshaw (2024) demonstrated that mutation bias can inhibit the runaway process in such cases.

#### How does the genetic basis of the male display affect the long-term outcome of the runaway process?

The above review of previous theories implies that the genetic architecture of the male sexual display can determine the long-term evolutionary outcome of the runaway process by affecting the mutation rate and mutation bias. The mutation rate of male sexual displays determines the likelihood of the loss of genetic variance and covariance. Mutation bias in male sexual displays can enable the stable maintenance of extreme phenotypes of the male display and female mate preference that have evolved through runaway.

We hypothesize that the runaway process can cause not only the evolution of an exaggerated male display and a biased female mate preference, but also their subsequent stable maintenance, provided that the male display is a complex trait governed by the omnigenic model (Box 1; Boyle et al. 2017; Liu et al. 2019). This is because omnigenic traits are expected to exhibit (1) a high mutation rate, (2) constraints on achievable phenotypes, and (3) strong mutation biases near the boundaries of the achievable phenotypic range. The high mutation rate is facilitated by the large mutational target size conferred by numerous peripheral genes (trans-regulatory regions)—a mechanism conceptually analogous to the genic capture hypothesis (Tomkins et al. 2004). Additionally, existing core genes impose upper and lower limits on the range of phenotypes that can evolve on microevolutionary timescales, which are primarily driven by mutations affecting the expression levels of core genes. Consequently, mutation bias is expected to arise in genotypes with phenotypes close to the boundary of this constrained phenotypic range. These three conditions are difficult to satisfy simultaneously when the male display is controlled solely by genetic loci with simple additive effects. Under additive genetic models with limited phenotypic ranges, increasing the number of loci inevitably reduces the phenotypic effect size of each individual locus; consequently, an expanded mutational target size fails to increase the supply of heritable phenotypic variation.

We examine the theoretical plausibility of this hypothesis using an individual-based evolutionary simulation model, in which the genetic architecture of a male display is determined by a set of parameters (Figure 4; for more details of the model, see Supplementary Appendix 1). The model assumes that the male display (*s*) is directly influenced by a fixed set of core genes, each of whose expression levels is regulated by an adjacent cis-regulatory region and many trans-regulatory regions located at random positions of the genome. As for female mate preference (*p*), we assume that it is controlled by a fixed number of loci with simple additive effects. For simplicity, we assume that the phenotypic evolution of the male display occurs exclusively through mutations in these regulatory regions. The model thus focuses on short-term evolutionary dynamics, during which the constitution of the core genes remains constant. When relatively small numbers of core genes are influenced by numerous trans-regulatory regions, the male display can be considered as an omnigenic trait (we hereafter refer to such situations as “omnigenic model”, although the original omnigenic model considers more complexities of gene regulatory networks, as we summarized in Box 1). In contrast, when there are many core genes (and their associated cis-regulatory regions) but no trans-regulatory regions, the genetic basis of the male display becomes equivalent to a conventional quantitative-genetic model with many additive loci.

**Figure 4.**
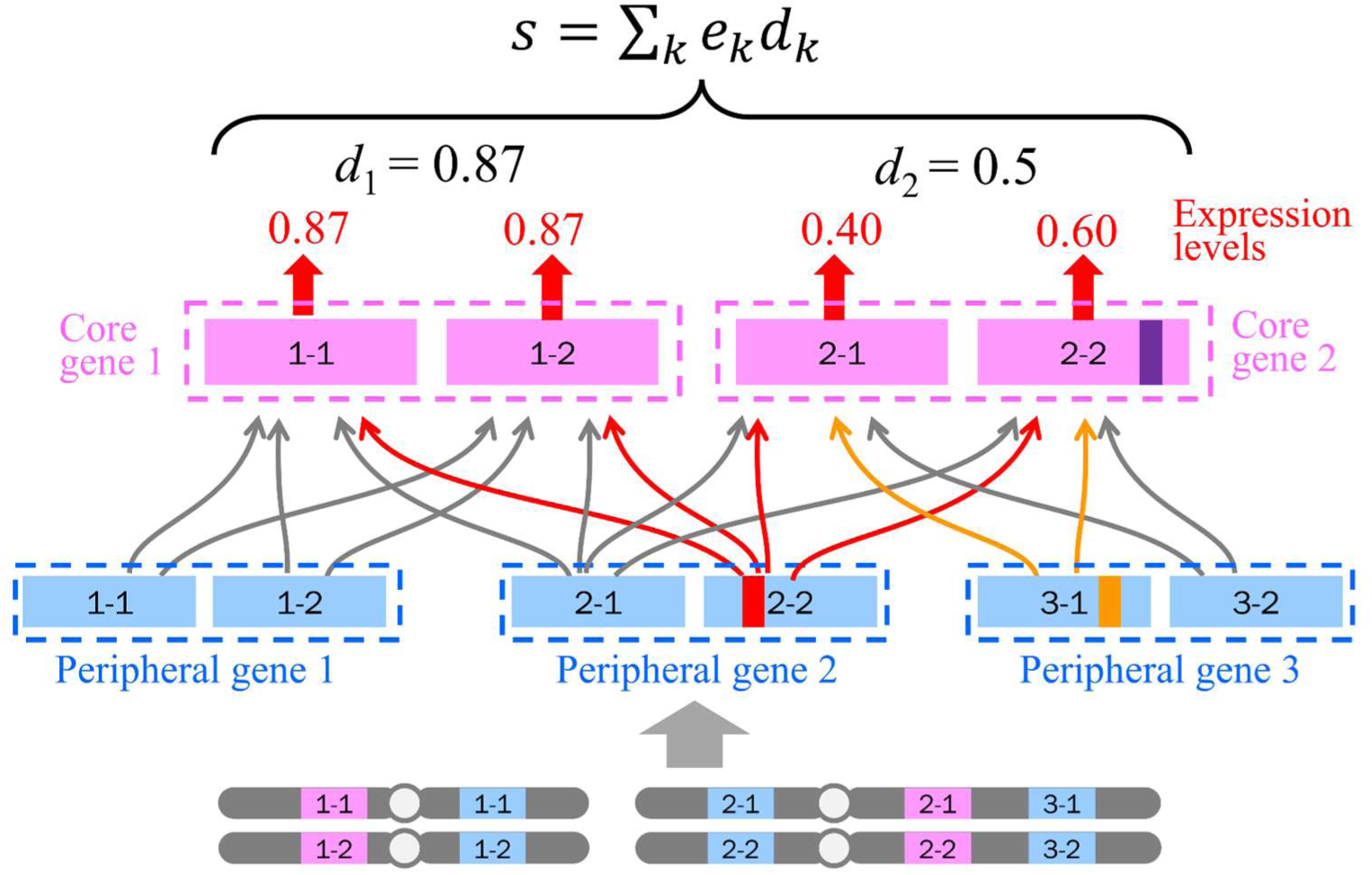
Schematic illustration of the genetic architecture of the male sexual display in our individual-based simulation model. The model assumes multiple core genes and numerous peripheral genes located at random positions in a genome consisting of 10 chromosomes of 2× 10^8^ bp long, which equals a total recombination distance of 200 cM. Core genes directly contribute to the developmental process of the male display, whereas peripheral genes produce transcription factors that can regulate the expression levels of core genes at any position in the genome. Each core gene is regulated by a cis-regulatory region located within the core gene and by a fixed number of peripheral genes (or trans-regulatory regions) via transcription factors. Thus, mutations in alleles of peripheral genes (red and yellow small squares) change the expression levels of all alleles of the core genes affected by them, whereas a mutation in the cis-regulatory region of an allele of a core gene affects only the expression level of that allele. The model adopts the “house-of-cards” approximation for the expression pattern of each core gene; any small change in the genetic sequence of any cis -or trans-regulatory region of a core gene changes its expression level within the range from 0 to 1. On the other hand, the model assumes that the process by which multiple core genes develop the phenotype of the sexual display follows a predictable manner: the value of *s* is simply determined by summing the phenotypic effect sizes of core genes weighted by their expression levels. The model also assumes that the functions of core genes themselves never change during simulations; thus, our simulations focus on microevolution driven by regulatory mutations on timescales where the constitution of core genes remains unchanged.

**Figure 5.**
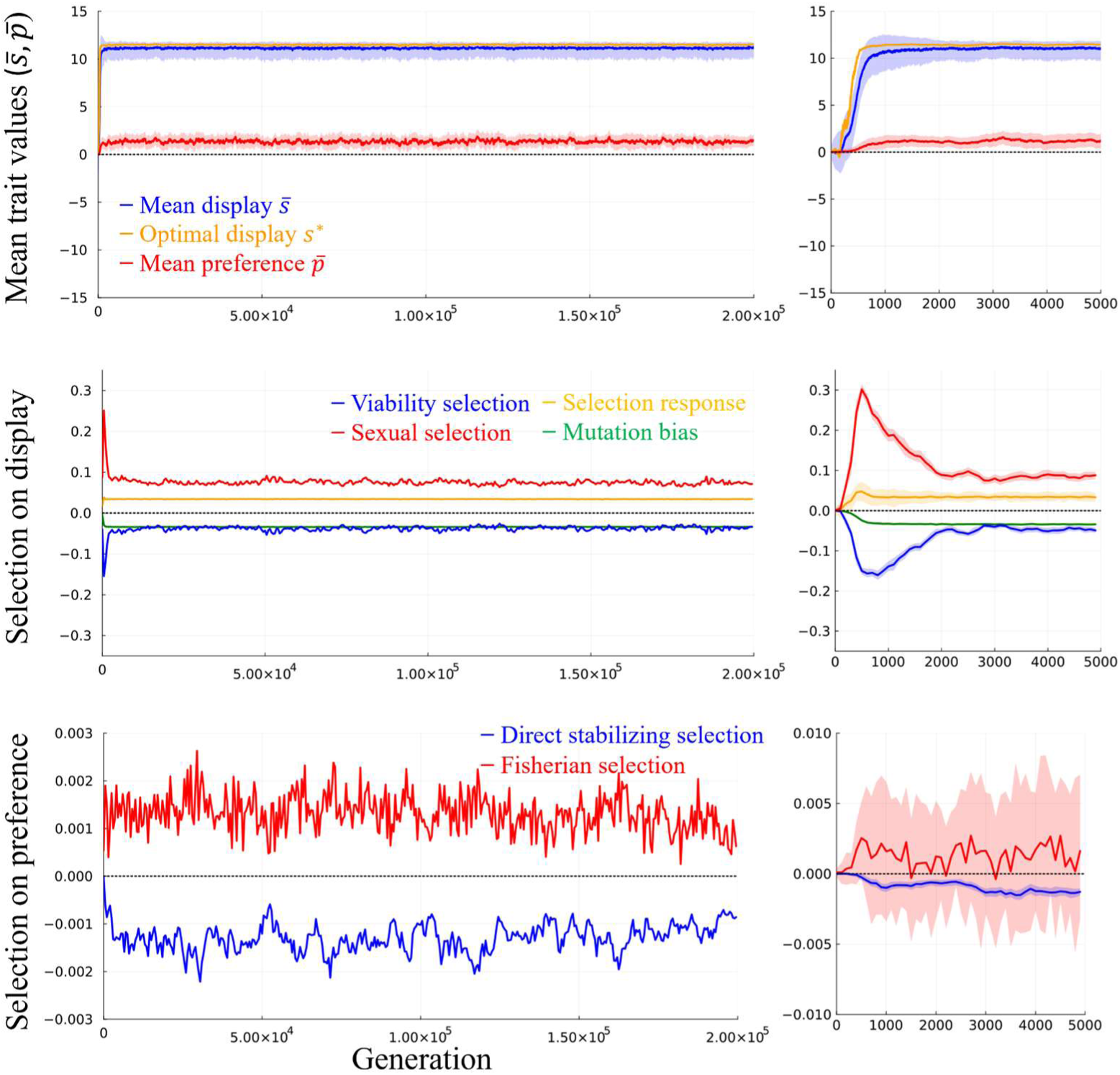
Example of evolutionary dynamics with an omnigenic male display. (a) Evolutionary dynamics of the population means of the male display (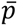; blue) and female mate preference (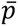; red). Colored areas show the ranges from the 5th to 95th percentiles for *s* and *p* (the ranges were very narrow in this panel). The yellow line shows the dynamics of the optimal male display value. (b) The ratio of the additive genetic covariance between the male display and female mate preference *B* to the additive genetic variance of the male display *G*. The dashed line shows the threshold value above which runaway starts in the classical quantitative genetic model (*B*/*G* ≥ 2*c*/*a*; Lande 1981). (c) Selection pressures on the male display. Blue and red lines show selection differentials on *s* in males caused by stabilizing natural selection before maturation and sexual selection after maturation, respectively. The yellow line shows the theoretically expected selection response in the absence of spontaneous mutations for each generation, which was determined using the mean phenotype calculated from modified genotypes in which all mutations introduced in the latest generation were removed, *s*’ (see Supplementary Appendix 2 for details). Similarly, we determined the effects of mutation bias for each generation (green line) by comparing genotypes with and without spontaneous mutations. (d) Selection pressures on the female mate preference. The blue line shows the selection differential on *p* in females caused by direct stabilizing selection. The red line shows the selection differential on *p* in males caused indirectly by sexual selection on the male display *s* combined with the genetic correlation (i.e. Fisherian selection). Values of *B*/*G*, selection differentials, selection response, and mutation bias were grouped into 100-generation bins, with the mean values displayed for each bin. Colored areas show the ranges from the 25th to 75th percentiles of selection differentials within the bins. Parameters: *N* = 40; *N* = 160; *N* = 2000; *M* = ∞; *θ* = 0.01; *K* = 5000. Other parameters were set to their default values (Table 1).

We tailor our model assumptions to suit our objective of investigating the runaway process. First, the model considers only Fisherian selection and direct costs of extreme sexual displays and biased mate preferences. Other selection pressures, such as good-gene effect, sexual conflict, and direct benefits of mate choice are not incorporated. Since the model considers direct stabilizing selection on female mate preference, the combination of random mating and the least costly male display is evolutionarily stable when runaway does not occur (Supplementary Figure S1). Second, we model female mate preference using an exponential function as in many previous models of sexual selection, including Lande’s (1981) psychophysical model. This modeling approach introduces severe artificial constraints on the mate preferences that can evolve; nevertheless, we adopt this approach for three reasons. First, evolutionary dynamics under this model are relatively well-understood and comparable with previous theoretical results. Second, runaway can occur under a wide range of conditions when the preference function is the exponential type, making it suitable for studying the runaway process. Third, this modeling approach allows a single quantitative-genetic trait to determine the entire shape of the preference function, which simplifies analysis and interpretation of simulation results. Although the biological validity of this assumption is questionable, we adopted this simple assumption that favors the occurrence of the runaway process in order to investigate how the runaway process may halt.

#### Simulation results

Using this model, we simulated evolutionary dynamics under systematically varied genetic architectures for the male sexual display by varying the number of core genes and their cis-regulatory regions (*N*) and the number of trans-regulatory pathways controlling each core gene (*N*), while keeping the number of peripheral genes (*N*) constant (20 ≤ *N* ≤ 320; 10 ≤ *N* ≤ 200; *N* = 2000). We also assumed that all the core genes have the same phenotypic effect size (*ε*) if their expression levels are identical. Therefore, the larger *N*, the larger the upper and lower limits of the male display phenotype that can evolve. Additionally, we compared simulations under two alternative genetic architectures of the male display: “additive” (20 ≤ *N* ≤ 320; *N* = *N* = 0) and “omnigenic” (20 ≤ *N* ≤ 320; *N* = 160; *N* = 2000). With these genetic architectures, we conducted simulations under various intensities of sexual selection by varying the upper limit on the number of matings per male (*M*). When single males can mate with many females, strong sexual selection can occur since the disparity in mating success among males can become large. Conversely, sexual selection becomes very weak when every male can mate with only a single female (*M* = 1; i.e., monogamy). We conducted simulations with *M* ≥ 2.

Simulations showed that the long-term evolutionary outcomes of Fisherian runaway vary depending on the genetic architecture of the male sexual display. We identified four mechanisms by which the runaway process was halted in the simulations: (1) evolutionary suicide, (2) the loss of selection gradient, (3) the loss of genetic variance (lek paradox), and (4) stable maintenance of an exaggerated sexual display. Methods used to categorize simulation results are provided in Supplementary Appendix 3.

Evolutionary suicide occurred under the combination of large values of *N* and extremely large values of *M*, irrespective of the genetic model (Figure 6). Under such conditions, there is virtually no limit to the male display value that can evolve, and the intensity of sexual selection never saturates with an increase in female mate preference. Hence, once a runaway process started, a continuous increase in the intensity of the male display and female preference led to a monotonic increase in the male survival cost of bearing exaggerated displays, resulting in evolutionary suicide. Figure 3a shows an example. While the initial population consisted of females without a biased mate preference and males without a costly sexual display, a rapid evolution of mate preference and sexual display occurred in the period between generation 1300 and 1500. In this period, the evolution of female mate preference caused the optimal trait value of the male display to move faster than the evolution of the male display itself, indicating the occurrence of runaway. The evolution of an exaggerated male display decreased the survival rate of males and the genetic variation in male displays. The population eventually went extinct due to the accidental mortality of all males.

**Figure 6.**
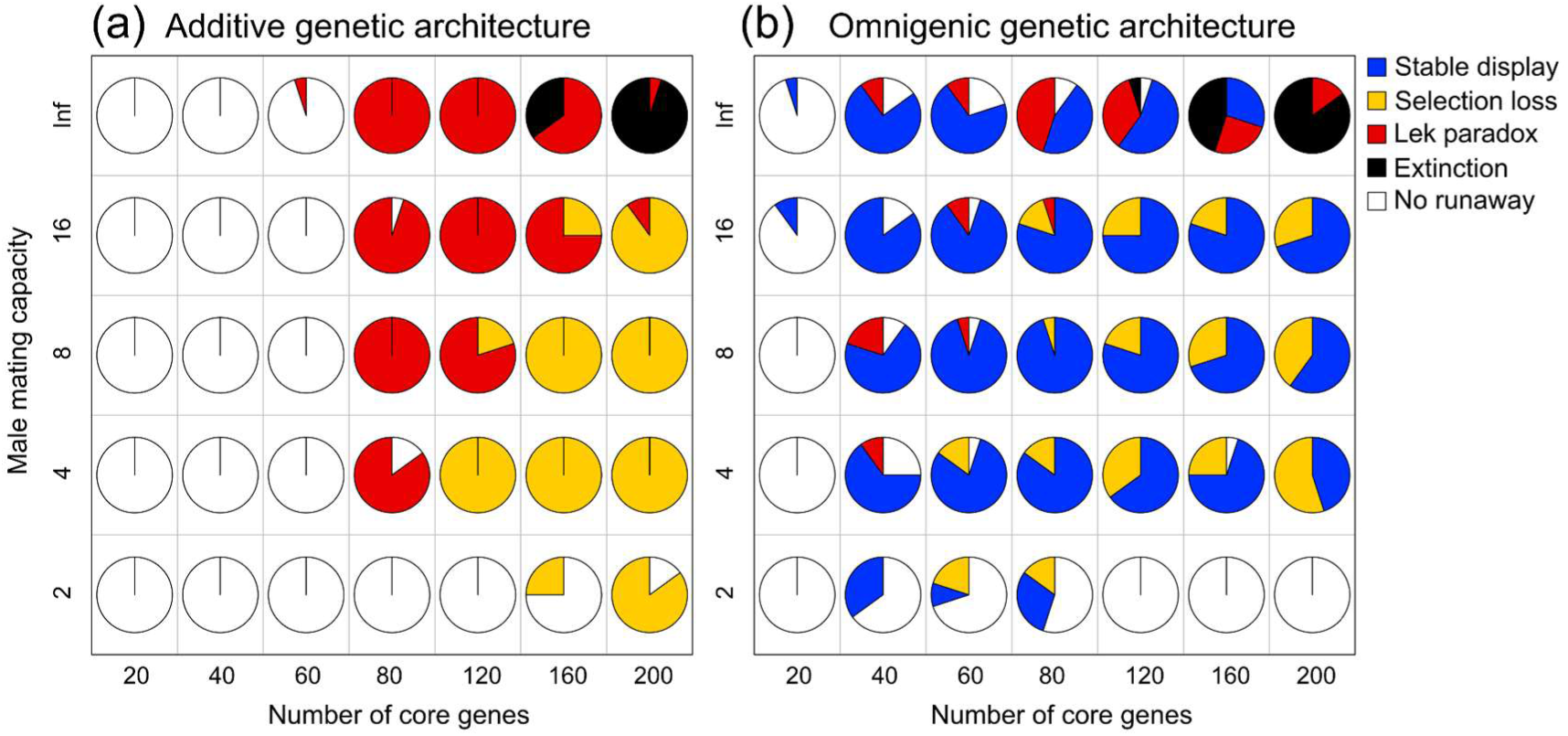
Effects of the maximum male mating capacity (*M*; vertical axis), which controls the intensity of sexual selection, and the number of core genes (*N*; horizontal axis), which sets limits on the phenotypes that can evolve. Each pie chart shows the frequencies of the following evolutionary outcomes of runaway across 20 simulation replicates under the same parameter conditions: stable maintenance of an exaggerated male display (blue), cyclic evolution due to the loss of the selection gradient (yellow), cyclic evolution due to the loss of genetic variation (red), evolutionary suicide (black), and failure of runaway (white). (a) Results of simulations with a male display controlled solely by loci with simple additive effects (Supplementary Figure S2 provides results with larger *N*). (b) Results of simulations with an omnigenic male display controlled by *N* core genes, each of which is regulated through 160 pathways from a total of 2,000 peripheral genes. Other parameters were set to their default values (Table 1).

The loss of the selection gradient on the male display was a common evolutionary outcome under the additive genetic model with the combination of large values of *N* and small values of *M* (Figure 6a; results with larger *N* are in Supplementary Figure S2). With large values of *N*, the runaway process is not halted by the constraint on the male phenotypic range. However, with small values of *M*, the intensity of sexual selection saturates to a certain level under strong mate preferences. Consequently, runaway stops when the intensity of sexual selection is balanced out by the stabilizing natural selection on the male display. Figure 2e shows an example. The runaway process ceased when the average male display value (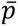; blue line) reached the optimal value (*s*^∗^; yellow line), at which the selection gradient on the male display vanished. This also led to the loss of indirect selection on female mate preference (Fisherian selection). As detailed above (see “(B) The loss of selection gradient can halt runaway”), the loss of Fisherian selection, combined with weak costs of mate choice, triggered a rapid evolution resulting in the loss of both the male display and female preference. Subsequently, the runaway process started again, leading to cyclic evolutionary dynamics in the long term.

The loss of male genetic variation (lek paradox) occurred under the additive genetic model with the combination of moderate values of *N* and large values of *M* (Figure 6a). Under such conditions, the evolution of a highly costly male display via runaway led to the reduction of the effective population size of males, which in turn reduced the genetic variance of the male display. Then, the paucity of male genetic variation weakened Fisherian selection through two pathways. First, the genetic uniformity among males eliminated the fitness disparity among them, which is necessary for Fisherian selection to operate. Second, without genetic variation among males, female mate choice did not generate genetic covariance between the male display and female preference, another requirement for Fisherian selection. Figure 3b shows an example. This simulation assumed a male display controlled only by loci with simple additive effects. Initially, the runaway process drove the evolution of a biased female mate preference and an exaggerated male display. In this period, the additive genetic variance of the male display increased, which is consistent with the argument of Xu & Servedio (2026) that the runaway process can increase the genetic variance of the sexual display. However, the evolution of a highly costly sexual display inevitably led to the reduction of the effective population size of males, which caused the subsequent loss of the additive genetic variance of the male display. Consequently, the runaway process ceased due to the lack of male genetic variation before the average male display (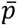; blue line) reached the optimal value (*s*^∗^; yellow line). As shown in Figure 3b, both sexual selection on males and Fisherian selection on females vanished when the runaway process ceased (methods to quantify Fisherian selection are provided in Appendix 2). The loss of Fisherian selection due to the lek paradox was followed by a rapid evolution resulting in the loss of the biased female mate preference and exaggerated male display by the same mechanism as in the case of the loss of selection gradient. Thus, cyclic evolutionary dynamics were observed in the long term.

Stable maintenance of an exaggerated male display (Figure 5) occurred when the male display was an omnigenic trait (Figure 7; 40 ≤ *N* ≤ 160; *N* ≥ 160; *N* = 2000). Moderate intensities of sexual selection (4 ≤ *M* ≤ 16) were most favorable for this evolutionary outcome (Figure 6b). Under such conditions, an exaggerated male display that was close to a boundary of the possible phenotypic range was stably maintained in an equilibrium state. This was because both the genetic variation of the male display and the selection gradient on it did not vanish at evolutionary equilibrium states, whereby Fisherian selection maintaining a biased female preference did not vanish either. The loss of the selection gradient did not occur because sexual selection was canceled not only by natural selection but also by mutation bias in such equilibrium states. We confirmed this by quantifying mutation bias and selection pressures in a simulation example (Figure 5c, d; Appendix 2). The loss of genetic variation (lek paradox) did not occur owing to high phenotypic mutation rates attained by numerous peripheral genes inflating the mutational target size.

**Figure 7.**
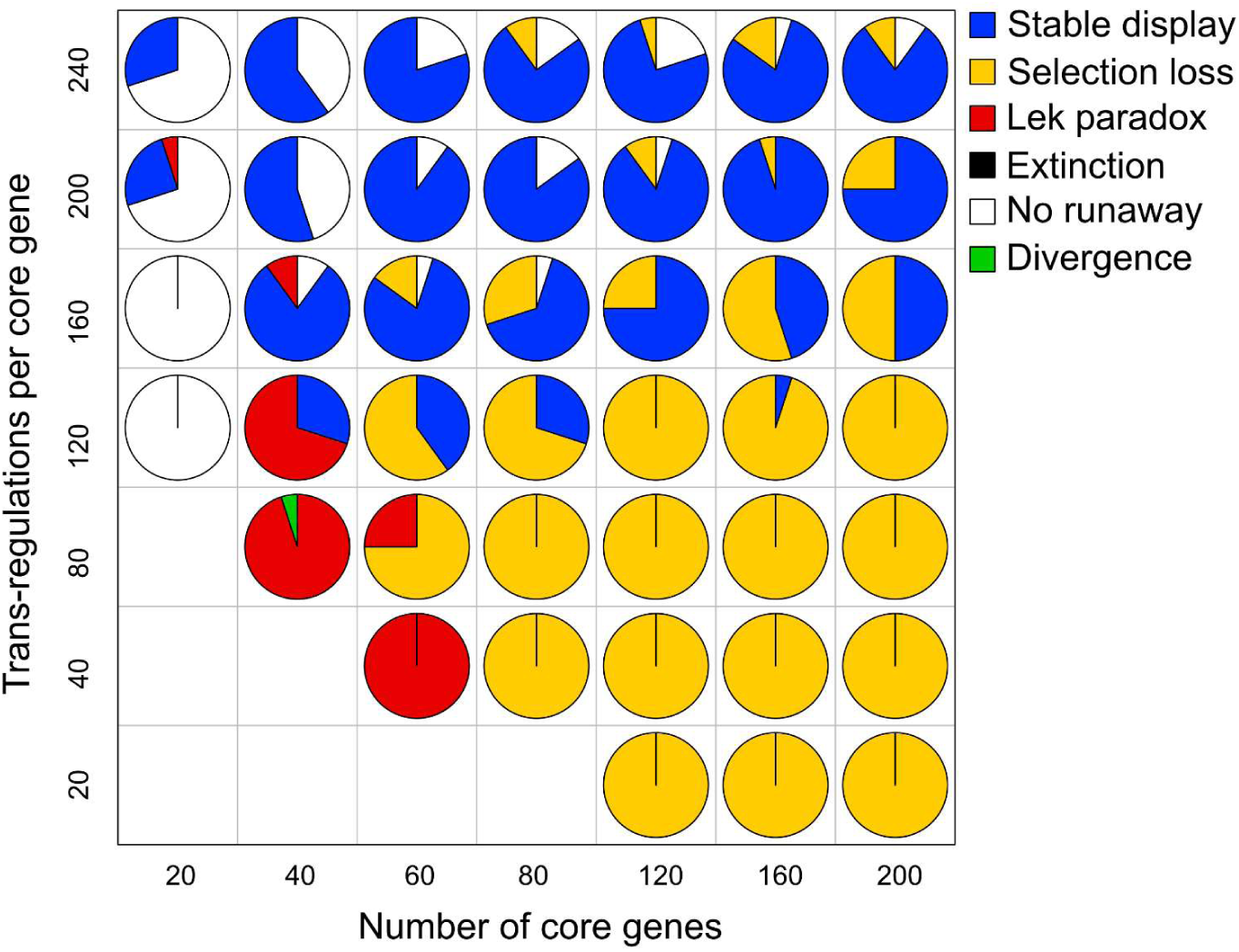
Effects of the number of core genes (*N*; horizontal axis) and the number of trans-regulatory pathways affecting each core gene (*N*; vertical axis). Each pie chart shows the frequencies of the following evolutionary outcomes of runaway across 20 simulation replicates under the same parameter conditions: stable maintenance of an exaggerated male display (blue), cyclic evolution due to the loss of the selection gradient (yellow), cyclic evolution due to the loss of genetic variation (red), failure of runaway (white), and ephemeral sympatric divergence (green). Parameter combinations with no pie chart violated the condition *N* × *N* ≥ *N*, which was postulated in our simulation model (see Methods). Other parameters were set to their default values (Table 1).

#### Model implications

These results demonstrate that the genetic architecture of the sexual display can govern long-term evolutionary dynamics with Fisherian runaway under non-ideal conditions where the population size is finite, males cannot mate with infinitely many females, and the genetic variance-covariance matrix is not fixed. When the male display was controlled by loci with simple additive effects as in the classical additive infinitesimal model, the dominant long-term evolutionary outcome of runaway was cyclic evolutionary dynamics. Such dynamics occurred because the runaway process halted either by the loss of the selection gradient on the male display or the loss of additive genetic variance of the male display (lek paradox). The loss of the selection gradient occurred when constraints on male mating capacity limited the intensity of sexual selection. When sexual selection was allowed to become intense, the runaway process inevitably led to a reduction in the effective population size of males, causing the loss of the additive genetic variance of the male sexual display.

In contrast, when the sexual display was an omnigenic trait, an exaggerated sexual display and a biased mate preference that have evolved through runaway were able to persist stably for the long term under a wide range of conditions. In this case, the lek paradox did not occur because numerous peripheral genes facilitated the maintenance of male genetic variation. Simultaneously, the loss of the selection gradient was avoided with mutation bias toward less extreme sexual displays (Iwasa & Pomiankowski 1995), which occurred because there were upper and lower limits to phenotypes that could evolve with a fixed set of core genes. These effects together enabled the stable maintenance of exaggerated sexual displays. It would generally be expected that existing functional genes set limits on phenotypes that can evolve through regulatory mutations alone. Under such conditions, regulatory mutations in genotypes with extreme phenotypes should tend to generate mutation biases toward less extreme phenotypes. Therefore, the stable maintenance of exaggerated sexual displays via mutation bias (Iwasa & Pomiankowski 1995) may be more feasible than previously thought when phenotypic evolution is mainly driven by regulatory mutations.

It is also worth noting that Fisherian selection (indirect selection on female mate preference) tended to exhibit substantial temporal fluctuation even at evolutionary steady states. Accordingly, although Fisherian selection could promote the maintenance of biased female mate preferences, this effect could hardly be detected without measuring the genotypes and fitness of all individuals in the population over many generations. These results imply that empirical studies failing to detect Fisherian selection do not necessarily invalidate the hypothesis that sexual displays and female preferences in the current population had evolved through Fisherian selection in the past.

#### Remaining issues

As in many existing models of sexual selection, only open-ended shapes of the mate preference function could evolve in our model using exponential function to link genotypes and the shape of the preference function. Although this assumption facilitated the comparison between our model and existing theoretical models, the evolution of preference functions in real organisms is unlikely to be subject to such a strong constraint, and evolutionary dynamics can change depending on how preference functions are formulated (Fry 2024; Kagawa 2026; Xu *et al*. 2023). It has been shown that Fisherian runaway is more likely to occur when a mate preference is constrained to open-ended preferences than when it is constrained to unimodal preferences (Hall *et al*. 2000; Lande 1981). Additionally, Fry (2024) suggested that relative mate preferences are more favorable for runaway than absolute mate preferences. Since runaway occurs only under limited model assumptions, the feasibility of runaway in real organisms remains controversial (Fry 2024). However, mutations in real organisms may generate diverse mate preferences including both unimodal and open-ended preferences, and runaway may start when an open-ended and relative preference is generated by a mutation, even if such mutations are rare. Thus, although the likelihood of runaway may be lower in real organisms than in our model, we suggest that runaway generally makes random mating evolutionarily unstable when mutations can generate diverse mate preferences.

Since our model considered only open-ended mate preferences, it underestimates the variation of evolutionary dynamics that runaway can generate. For example, the runaway process can promote the evolution of mate preferences for rarer male phenotypes when the evolution of frequency-dependent mate preferences is allowed (Kokko *et al*. 2007; Potter *et al*. 2023). Currently, little is known about the evolutionary dynamics that may arise through runaway when mutations can generate various types of mate preferences and mating systems.

While our model considered a range of genetic models including the omnigenic model, it still simplified the gene regulatory network by shrinking it into a network with only two layers (see Box 1 and methods). Also, the model incorporated neither gene-environment interactions nor deleterious mutations, while they can be major sources of heritable phenotypic variation of the male display and good-gene benefits of mate choice (Dugand *et al*. 2019; Kokko & Heubel 2008; Rowe & Houle 1997; Tomkins *et al*. 2004). In the presence of good-gene effects, sharing of genetic pathways between sexual displays and fitness-relevant traits can promote the evolutionary stability of honest signaling of male condition (Roulin 2016; Weaver *et al*. 2017). It has also been suggested that sexual displays that have originally spread through the runaway process can evolve into indicator traits of good genes when there are pleiotropic mutations affecting both displays and viability (Chandler *et al*. 2013). Future theoretical studies should clarify how gene regulatory networks underlying sexual displays affect evolutionary dynamics under sexual selection with both sexy-son and good-gene benefits of mate choice.

More generally, the evolution and maintenance of female mate preferences can be promoted by various other mechanisms including good-gene and direct benefits of mate choice, sexual conflict, reinforcement selection, and ecological selection on traits affecting mate preference (Kokko *et al*. 2006; Kuijper *et al*. 2012; Rosenthal 2017). They can operate together with Fisherian selection, which occurs whenever both female mate preference and male sexual displays are heritable. An important remaining issue is how the genetic architecture of sexual displays affects long-term evolutionary dynamics with multiple types of selection pressure on mate preference.

## Conclusions

Based on a review of existing theories and individual-based simulations, we categorized the mechanisms that halt the runaway process into four types. The categorization is based on which of the idealized assumptions of Lande’s (1981) model become a poor approximation as a result of evolutionary changes caused by the runaway process: an infinitely large population size, a constant additive genetic variance-covariance matrix, an unlimited phenotypic range, and unlimited male mating capacity. Runaway can halt when it leads to population extinction (evolutionary suicide; category A), the loss of the selection gradient on the male sexual display (category B), or the loss of genetic variation of the male display (lek paradox; category C). In categories B and C, runaway drives cyclic evolutionary dynamics in which the evolution and loss of a biased female mate preference and an exaggerated male display repeat. Additionally, the runaway process can halt when the male display approaches a boundary of the range of phenotypes that can evolve (category D). In this case, exaggerated male displays can stably persist in equilibrium states where sexual selection is balanced by stabilizing natural selection plus mutation bias.

Our individual-based simulations revealed key conditions for the four categories of long-term outcomes of Fisherian runaway under non-ideal conditions. An important finding is that the genetic architecture of the sexual display determines whether an exaggerated sexual display and a biased mate preference that evolved through runaway can persist stably in the long term. With simple additive genetic architectures, there were no mechanism to prevent the loss of the selection gradient (category B) and the lek paradox (category C). With omnigenic architectures, in contrast, both high phenotypic mutation rates and mutation biases toward less exaggerated phenotypes were attained through key features of the gene regulatory network, which prevented the lek paradox and the loss of selection gradient. These results show that the idealized assumptions in classical models of Fisherian runaway exert profound influences on long-term evolutionary dynamics, qualitatively changing the ultimate outcomes of the runaway process. Thus, although simplified models are highly useful for understanding evolutionary mechanisms, our findings underscore that models with biologically realistic assumptions are also crucial for interpreting empirically observed evolutionary dynamics driven by sexual selection.

## Box 1. The omnigenic model

Growing genomic evidence suggests that many complex traits align with the omnigenic inheritance model (Boyle *et al*. 2017; García-González & O’Reilly 2026; Liu *et al*. 2019). Unlike the classical additive infinitesimal model, this model classifies genes into two categories: core genes, which are directly involved in the biological pathways underlying a focal trait, and peripheral genes, which indirectly affect the trait by regulating the expression levels of the core genes. Generally, gene regulatory networks within cells are densely interconnected so that the effects of each gene can propagate throughout the entire network. Consequently, the expression levels of core genes can be influenced by nearly all genes that are actively transcribed within the same tissue or cell type. Based on this premise, the omnigenic model assumes that many complex traits are determined by the expression levels of a relatively small number (e.g., tens) of core genes that are affected by numerous (e.g., thousands) peripheral genes (Boyle *et al*. 2017; Liu *et al*. 2019). This theory was proposed to explain the "missing heritability" problem—the fact that for many complex traits including common diseases and human height, alleles significantly associated with phenotypes in genome-wide association studies (GWAS) account for only a small fraction of the observed heritability. Furthermore, the apparent paradox that many risk variants for common diseases lack clear biological links to the diseases based on known gene ontologies can be resolved if they are peripheral genes (Boyle *et al*. 2017). To date, the validity of the omnigenic model has been demonstrated across multiple complex traits (García-González & O’Reilly 2026; Ružičková *et al*. 2024; Sinnott-Armstrong *et al*. 2021; Vuckovic *et al*. 2020).

The individual-based model of the present study considers the evolution of an omnigenic trait with spontaneous mutations. The model considers tens to hundreds of core genes, whose gene expression levels are affected by thousands of peripheral genes. Owing to the presence of numerous peripheral genes, the phenotypic mutation rate reaches the order of 10^-2^ per generation, which is within the range of typical values for complex traits (Lynch & Walsh 1998), given a realistic mutation rate (from 10^-6^ to 10^-5^ per locus). While our model divides genes into core and peripheral genes, it simplifies gene regulatory networks by treating direct and indirect regulation without distinguishing between them. More specifically, our model considers a gene regulatory network with two layers: the upstream layer consisting of peripheral genes and the cis-regulatory regions of the core genes, and the downstream layer consisting of core genes, whose expression levels are regulated by genes in the upstream layer. Additionally, the model assumes that expression level of each core gene is randomly changed by any single mutation in any regulatory region that affects the core gene, instead of tracking gene expression dynamics within cells. This approximation is based on the assumption that indirect regulatory effects on core genes exerted by peripheral genetic variants are highly unpredictable due to the complexity of gene regulatory networks through which these effects propagate.

## Box 2. Lande-Kirkpatrick process

The models of Lande (1981) and Kirkpatrick (1982) demonstrated not only that Fisherian selection can cause runaway under certain conditions but also that evolutionary dynamics driven by Fisherian selection can have infinitely many neutrally stable equilibrium points under other conditions. The latter situation is often referred to as a “walk-towards” (Mead & Arnold 2004). The “Lande-Kirkpatrick process” (Prum 2010) refers to evolutionary dynamics driven by Fisherian selection, encompassing cases both where runaway occurs and where it does not.

In Lande’s (1981) model with “psychophysical preference”, runaway occurs when *B*/*G* > 2*c*/*a*, as described in the main text. When *B*/*G* < 2*c*/*a*, on the other hand, all equilibrium points on the line 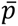 = 2*c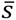*⁄*a* become neutrally stable. For example, when *B* = 0 (Figure 1a), female mate preference 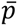 does not change over time as d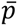/d*t* = 0, while the male display evolves to an equilibrium point (s*) at which sexual selection is balanced by the stabilizing natural selection, resulting in *W* (s*) = 0. Such a stable equilibrium point exists for every value of 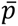. Although directional phenotypic evolution halts once a population reaches one of the stable equilibrium points, genetic drift can move the mean phenotype of a population (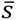, 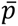) along the line of neutrally stable equilibria. This situation does not change qualitatively as long as *B*/*G*—the slope of evolutionary trajectory returning to the equilibrium line— is smaller than 2*c*/*a*—the slope of the equilibrium line (Figure 1b). While Fisherian selection is not essential for the occurrence of neutrally stable equilibria, it can accelerate the random evolution along the line of neutrally stable equilibria (Tazzyman & Iwasa 2010).

The occurrence of neutrally stable equilibrium points depends on the strong assumption that female mate preference receives no direct selection (Kirkpatrick 1985). When female mate preference *p* is subject to direct selection, the selection gradient of *p* can become zero only at the peaks of the fitness landscape of direct selection. Thus, the line of neutrally stable equilibria shrinks to a set of stable equilibrium points at locally optimal values of *p*. In this sense, random drift along the line of neutrally stable equilibria is not a robust explanation for the evolution and maintenance of biased female mate preferences. A remarkable exception is the case where sampling acceptable mate candidates is costly but evaluating individual males is not. In such cases, the cost of having mate preferences vanishes once an acceptable male display is fixed in the population, making female mate preferences selectively neutral at the equilibrium state (Kokko *et al*. 2015; Waffender & Henshaw 2023).

## Box 3. Definition and criteria of runaway

We adopt the definition of runaway as the evolutionary process in which Fisherian selection leads to a self-reinforcing evolution of mate preferences. Although the distinction between evolutionary dynamics with and without runaway is clear in the simple models of Lande (1981) and Kirkpatrick (1982), it is generally difficult to make this distinction in more complex systems for several reasons. First, no biologically feasible evolutionary trajectory can be simply divided into runaway and non-runaway, because the runaway process cannot continue forever in reality; evolutionary trajectories with runaway must also contain periods without runaway. Second, in finite populations, the runaway process is affected by stochasticity. Since it is fundamentally impossible to rigorously differentiate the effects of deterministic evolutionary forces from those of genetic drift in observable allele frequency dynamics, the identification of a runaway process inevitably involves uncertainty. Furthermore, the evolution of the G-matrix caused by genetic drift alone can reverse the inequality *B*/*G* > 2*c*/*a*—the condition for runaway in Lande’s (1981) model—during evolution; thus, even in simulation models, whether runaway occurs cannot be determined by parameters alone when stochasticity is considered.

To our knowledge, there is no consensus on criteria to identify periods of runaway in observed evolutionary dynamics. Although it is difficult to draw a clear distinction between periods with and without runaway, evolutionary dynamics driven by the runaway process are expected to show certain characteristics that are measurable at least in simulations. First, the additive genetic covariance between the male display and female mate preference relative to the additive genetic variance of the male display should be high during the runaway process. Second, the self-reinforcing evolution of mate preferences via runaway causes the optimal value of the male display to move in a constant direction at a faster rate than the evolution of the male display itself. The optimal male display refers to the value of the sexual display *s* at which sexual selection is canceled out by stabilizing natural selection. Runaway can thus be detected as periods during which the evolution of the male display is chasing this moving optimal value. Third, runaway renders the least costly sexual display and random mating evolutionarily unstable, even in the presence of stabilizing natural selection in favor of them. Thus, the evolutionary instability of these traits can be considered as a signature of the occurrence of runaway.

## Ethics

This work did not require ethical approval from a human subject or animal welfare committee.

## Data accessibility

All Julia source code associated with this study will be deposited in a public repository upon publication of the manuscript.

## Authors’ contributions

K.K.: conceptualization, analysis, investigation, methodology, writing— original draft. K.K. & M.K: writing—review and editing.

## Declaration of AI use

AI-assisted technologies were used to identify typos and grammatical errors.

## Conflict of interest declaration

We declare we have no competing interests.

## Funding

This study was supported by the JSPS Research Fellowships for Young Scientists (20J00583), and Japan Society for the Promotion of Science KAKENHI grants (22K15163 and 25K09750).

## Acknowledgments

We also thank members of Kawata laboratory of the Graduate School of Life Sciences of the Tohoku University and Kitano laboratory of the National Institute of Genetics for help in the laboratory. A large part of simulations in this study was carried out under the ISM Cooperative Research Program (2025-ISMCRP-0010; 2026-ISMCRP-0015) of the Institute of Statistical Mathematics.

## Appendix 1: individual-based simulation model

The individual-based model (implemented in Julia) simulates the evolution of the intensity and direction of female mate preference (*p*) and a quantitative male display trait (*s*) in sexually reproducing diploid organisms with non-overlapping generations. Both *s* and *p* can take positive or negative values. Positive (negative) values of *p* indicate a preference for males with positive (negative) *s*, and females with *p* = 0 mate randomly. We did not constrain *s* and *p* to positive values to prevent mutation bias from making random mating evolutionarily unstable; if *s* and *p* are constrained to positive values, any mutations affecting mate preference in individuals with *p* = 0 will increase *p*, and this mutation bias alone can disrupt the stable maintenance of random mating.

We assume direct stabilizing selection for both female preference *p* and male display *s*. Females with extreme preferences are subject to a weak search cost that reduces their fecundity by a factor of *exp* (−*θp*^2^), where *θ* indicates the intensity of the search cost. Males survive until reproduction with a probability of *exp* (−*cs*^2^), where *c* indicates the cost of bearing sexual display. The least costly male display is *s* = 0. The operational sex ratio becomes female-biased when males bear a non-zero sexual display *s* (either positive or negative).

A female’s preference for a male’s trait value *s* is given as exp(*αps*), where *α* is a parameter that controls how strongly the female mate choice is affected by the preference *p*. Thus, the female trait *p* determines both the intensity and direction of preference. In a mating period, each female examines *N* randomly chosen males (we set *N* = 100) and mates with the *i*^th^ male from them with a probability of *exp*(α*ps*_i_)/ ∑_j_ *exp*(α*ps*_j_). Thus, the preference is relative, and every female is guaranteed to mate with a male. The model also incorporates a weak cost of being choosy: female mate preference is subject to a weak stabilizing selection in favor of random mating (*p* = 0), which minimizes time and energy spent evaluating males. Therefore, strongly biased female mate preferences can evolve only when runaway makes random mating (*p* = 0) evolutionarily unstable.

The model assumes that the female trait is controlled by *L* loci with additive effects, whereas the male trait is controlled by *N* core genes, whose expression levels are affected by cis -and trans-regulatory regions. The phenotypic value of the male sexual display is given by 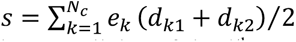, where *e* and *d* (*n* = 1, 2) are the effect size and expression levels of the two alleles of the *k*^th^ core gene, respectively. The model thus assumes that core genes have independent additive effects on the phenotypic value. The phenotypic value is none of the core genes are expressed. For simplicity, we assume that *e* does not evolve and that half of the core genes have *e* = 2*ε*, whereas the remaining half have *e* = −2*ε.* Additionally, we assume that the expression level *d* ranges from 0 to 1. Hence, the upper and lower limits of the male display phenotype are given by *s* = *εN* and *s* = −*εN*, respectively. The model assumes that *d* can change through genotypic evolution in two types of regulatory regions: cis -and trans-regulatory regions. We assume that each allele of core gene contains a single cis-regulatory region regulating its expression level; thus, the number of cis-regulatory regions equals the number of core genes, *N*. On the other hand, the number of peripheral genes (trans-regulatory regions) *N* can differ from that of core genes, and each peripheral gene can affect multiple core genes at any position in the genome. Unlike cis-regulatory regions, trans-regulatory regions affect alleles of their target core gene on both chromosome copies simultaneously. For simplicity, we assume that each core gene is regulated by a constant number (*N*) of peripheral genes. The topology of the regulatory network, which does not evolve, is determined before starting a simulation; for each of the *N* ×*N* trans-regulatory pathways, we randomly assigned a peripheral gene, while ensuring that every peripheral gene is involved in at least one regulatory pathway (to satisfy this, we assume *N* ×*N* ≥ *N*).

The model assumes that the expression level of each core gene (*d*) is highly sensitive to changes in genetic sequences of the regulatory regions affecting it. More specifically, randomly chosen values of *d*, ranging from 0 to 1, are assigned to every different genotype at all the regulatory regions for the core gene *k*. For simplicity, we assume no statistical correlations in the values of *d* among different regulatory genotypes. Hence, similar regulatory genotypes for a core gene do not necessarily have similar expression levels of that core gene; a single mutation in a regulatory region can entirely change the expression level of core genes affected by it. However, despite the highly unpredictable nature of mutational effects on the expression levels of single core genes, similar genome-wide genotypes tend to have similar phenotypes in this model. This is because all the core genes affect the phenotypic value only additively.

This model can reproduce three biological features that are expected to emerge in complex traits that follow the omnigenic inheritance model. First, there are upper and lower limits to the range of phenotypic values that can evolve, as explained above. This condition seems biologically plausible for short-term evolution mainly driven by regulatory changes. Second, mutation biases will occur for genotypes with extreme phenotypes that are close to the upper or lower limits. This is because finely tuned gene expression profiles are necessary to generate extreme phenotypes, whereas the vast majority of gene expression profiles generate moderate phenotypic values. For instance, when a genotype generates the upper-limit phenotypic value, the expression levels of all the core genes with positive and negative effects are 1 and 0, respectively; any random changes on this gene expression profile result in less extreme phenotypes. Third, the supply of heritable phenotypic variance per generation increases with the mutational target size (e.g., the number of loci). While this condition is biologically sound, it is not satisfied in most existing models that considered the evolution of polygenic traits with limited phenotypic ranges by assuming that the phenotypic effect size of each genetic locus is inversely proportional to the number of loci.

In this model, the genetic architecture of the male display can be changed by varying the values of three parameters: the number of core genes (*N*), the number of peripheral genes (*N*), and the number of trans-regulatory pathways for each core gene (*N*). When there are many core genes but no peripheral gene or trans-regulatory pathways (*N* = *N* = 0), the genetic basis of the male display becomes equivalent to that in a regular quantitative genetic model with many additive loci. When there are dozens of core genes regulated by many peripheral genes via numerous trans-regulatory pathways, the male display follows the omnigenic inheritance model. Mutations have epistatic effects more frequently when the number of trans-regulatory pathways (*N* × *N*) is large compared to the numbers of core and peripheral genes.

## Appendix 2: Methods to analyze evolutionary dynamics within single simulations

### Criteria of runaway

We determined whether runaway occurred during a given period in simulations based on three criteria (Box 3). First, we determined whether the additive genetic covariance between the male display and female mate preference (B) relative to the additive genetic variance of the male display (G) was high in the focal period. Although our model differed from Lande’s (1981) model in several important assumptions, the dynamics around the origin (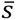 = 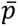 = 0) should be similar. Thus, the condition for runaway in Lande’s (1981) model—*B*/*G* > 2*c*/*a*—can detect periods during which the origin is evolutionarily unstable due to the runaway process. We estimated the additive genetic variance and covariance based on phenotypes of parents and offspring. Since the additive genetic variance of a trait is the product of the phenotypic variance and the heritability of the trait (*h*^2^), we estimated it by determining the heritability using linear regression of offspring phenotypes on midparent values. Similarly, we estimated the additive genetic covariance between two traits based on the covariance between midparent values of one trait and offspring phenotypes of the other.

Second, we detected periods during which the optimal male display value *s*^∗^ moved at a faster rate than the evolution of the male display itself. For this purpose, we first determined the optimal male display value (*s*^∗^) under fixed values of female mate preference by conducting evolutionary simulations without the evolution of female mate preference across a range of female mate preference values (Figure 2a). As expected, the slope of the equilibrium curve around the origin equals that of the equilibrium line in Lande’s (1981) model (1/2*c*). The equilibrium curve saturates at the male display value at which male survival rate becomes 1/*M* due to the costs of bearing the display. This is because male reproductive success cannot exceed the maximum number of mating partners per male, *M*. When the male survival rate is less than 1/*M*, every male mates with *M* females; sexual selection cannot increase in strength even if female mate preference (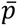) increases. Thus, the equilibrium curve saturates at *s* such that *exp*(−*cs ^2^*) = 1/*M*, which yields 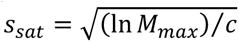. We then constructed a formula to estimate the optimal male display value *s*^∗^ as a function of 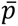 by fitting a cubic spline curve to the simulation results, using the *Spline1D* function from the *Dierckx* package in Julia. Using this formula, we calculated the time series of the optimal male display value (*s*^∗^) for our simulations to examine whether there were periods during which *s*^∗^ moved at a faster rate than the male display value 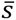.

Third, we incorporated stabilizing natural selection on both the male display and female mate preference so that the origin point (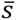 = 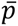 = 0) was evolutionarily stable unless runaway occurs. Therefore, evolutionary trends deviating from the origin indicate the occurrence of runaway in our model.

### Quantification of natural selection, sexual selection, Fisherian selection, and mutation bias

We quantified selection pressures in our simulations using selection differentials. In each generation, selection differentials caused by natural and sexual selection were calculated separately. The selection differential for the male display *s* due to natural selection is given by

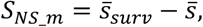

where 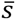 and 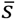_surv_ are the means of *s* among males before and after viability selection, respectively. The selection differential due to sexual selection is given by

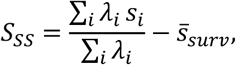

where *s* and *λ* are the sexual display and reproductive output of male individual *i*, respectively (*λ* = 0 for individuals that died before mating); the first term is the mean of *s* weighted by reproductive fitness. We measured Fisherian selection using the selection differential for the genotypic value of female mate preference (*p*) in males, which is given by

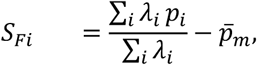

where *p*_i_ is the genotypic value of *p* of male individual *i*, and 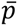_m_ is the mean of *p* among males before viability selection. *S*_Fisher_ accounts for indirect selection on *p* caused by natural and sexual selection on *s* in males. Similarly, we measured natural selection on female mate preference due to its costs using the following selection differential calculated in females:

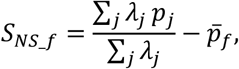

where *p* and *λ* are the mate preference and reproductive output of female individual *j*, respectively; 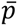_ƒ_ is the mean of *p* among females.

Note that the expected value of the selection response for female mate preference equals (*S*_Fisher_ + *S*_NS_ƒ_)⁄2, because the heritability of *p* is always 1 as *p* is determined solely by genetic loci with additive effects. On the other hand, the selection response for the male display is not necessarily proportional to the selection differential, because epistasis among alleles can affect the phenotypic value of *s*.

To quantify mutation bias for the male display, we calculated the phenotypic value of *s* from modified genotypes in which all mutations introduced in the latest generation were removed. Specifically, mutation bias was calculated as

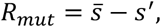

where *s*’ is the mean phenotypic value calculated across all genotypes in the population after removing newly introduced mutations. Additionally, to test whether the effect of selection on the male display is counterbalanced by mutation bias in stable equilibrium states, we calculated the theoretically expected value of the selection response in the absence of mutations as follows:

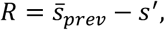

where 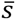_prev_ represents 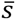 of the previous generation.

## Appendix 3: Methods to categorize simulation results

We categorized simulation results based on the observed dynamics of the population means of the male display (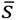) and female mate preference (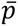), which were recorded every 100 generations during the period from generations 50,000 to 100,000 (records from the first 50,000 generations were excluded to eliminate the effects of the initial state). The first category is *evolutionary suicide*, in which the evolution of a highly costly male display led to population extinction. The second and third categories are *failure of runaway* and *sympatric divergence*, in both of which the distance of the mean female preference from 0 (|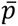|) was on average smaller than its standard deviation in the focal period (generations 50,000–100,000). This condition could be satisfied either when biased female mate preferences did not evolve (i.e., failure of runaway) or when female mate preference (*p*) diverged into positive and negative values within a population while keeping the mean female preference near 0 (i.e., sympatric divergence) (Figure S3). In the latter case, the male sexual display also diverged into two extreme phenotypes, increasing the cost of bearing sexual displays. Simulations satisfying both of the following conditions were categorized as sympatric divergence: (A) the male survival rate was on average lower than 0.8 in the focal period; (B) there was a period during which the phenotypic distribution of female mate preference (*p*) was bimodal. We checked condition (B) using the bimodality coefficient given by *BC* = (*g* ^2^+1)/(*g* ^2^+3), where *g* is the sample skewness and *g* is the sample excess kurtosis; the condition was satisfied if the average *BC* within the subset of time points representing the highest 10% of *BC* values was greater than 0.7. The fourth category is *stable maintenance of an exaggerated sexual display*, in which a combination of an exaggerated male display and a biased female mate preference that evolved through runaway remained virtually unchanged for a prolonged period: specifically, (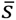, 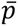) remained in the first quadrant of the (*s*, *p*) space during the focal period. In the remaining cases, we observed cyclic evolutionary dynamics in which the evolution to gain and lose an exaggerated male display was repeated. Such cases were classified as either *the loss of the selection gradient* or *the loss of genetic variation* based on the dynamics of the optimal male display value (*s*^∗^), which was estimated from the value of 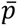 (Appendix 2). To detect the loss of the selection gradient, we first extracted a subset of time points corresponding to the highest 10% of the absolute values of the mean male display (|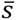|) from the entire time series in the focal period. Then, a simulation was categorized as *the loss of the selection gradient* when the value of |*s*^∗^ − 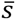| was on average smaller than the standard deviation of *s* within this subset of time points. When this condition was satisfied, the male display evolved to approach the optimal value, which caused the selection gradient on the male display to approach zero. Simulations with cyclic evolutionary dynamics that did not satisfy the above condition were categorized as *the loss of genetic variation*. In this case, the evolution of the male display ceased before 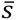 approached the optimal value *s*^∗^ (this often occurred near the upper or lower limit of possible phenotypes), and cyclic evolution occurred because Fisherian selection to maintain female mate preference became weak due to the reduction of the additive genetic variance of the male display.

---

**Figure S1.**
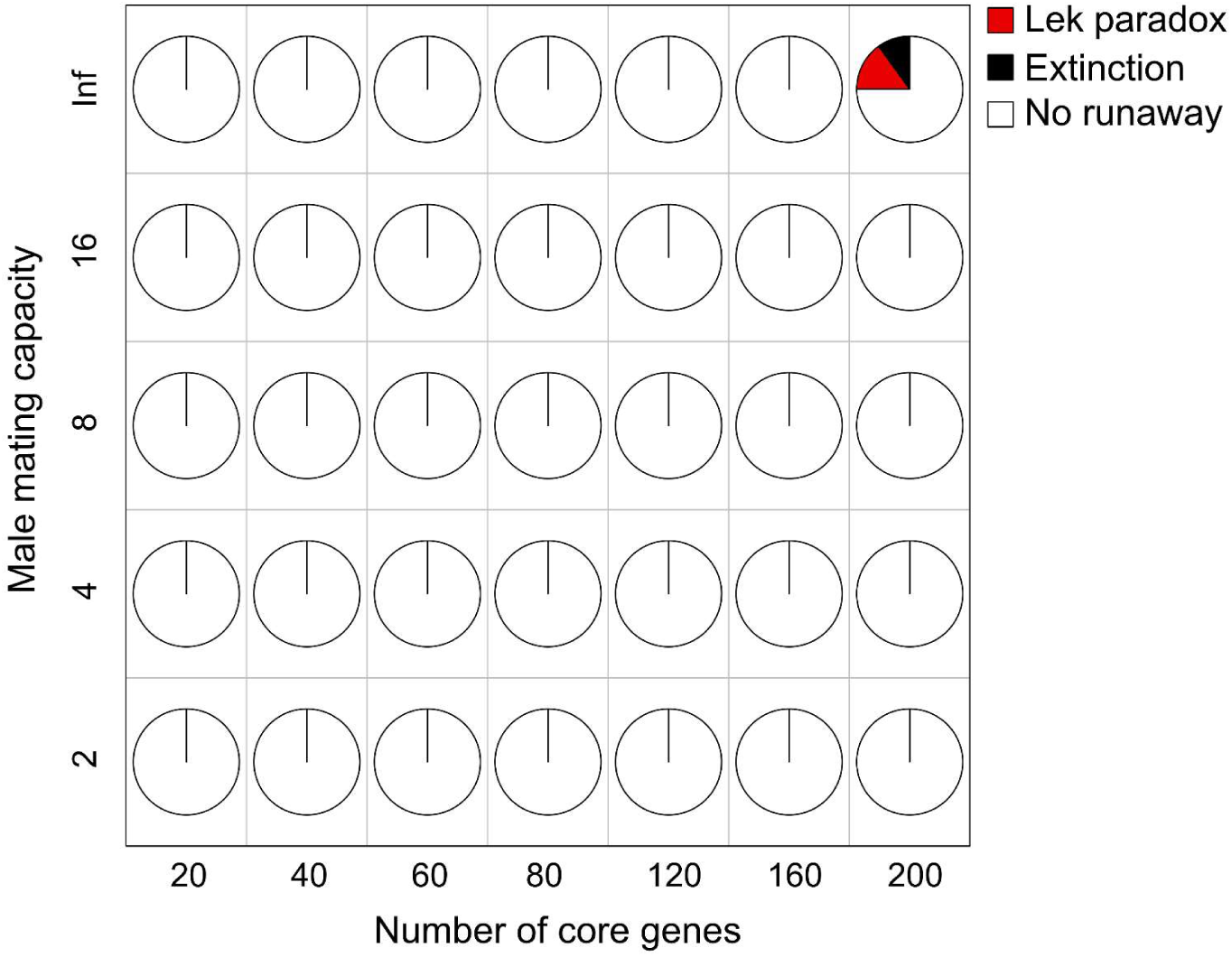
The runaway process was essential for the evolution of exaggerated sexual displays. In the simulations shown in this figure, the runaway process was artificially suppressed by randomizing mating partners without changing the intensity of sexual selection on males. Namely, after female mate choice, the mating partner of each female was swapped with the partner of another randomly selected female. Through this procedure, the effect of mate choice in generating genetic covariance between the male sexual display and female mate preference was suppressed, while the mating success of each male remained unchanged. Other simulation conditions including parameter values were the same as in the simulations shown in Figure 6b. Each pie chart shows the frequencies of the following evolutionary outcomes across 20 simulation replicates under each parameter condition: cyclic evolution due to the loss of genetic variation (red), evolutionary suicide (black), failure of runaway (white). Other parameters were set to their default values (Table 1).

**Figure S2.**
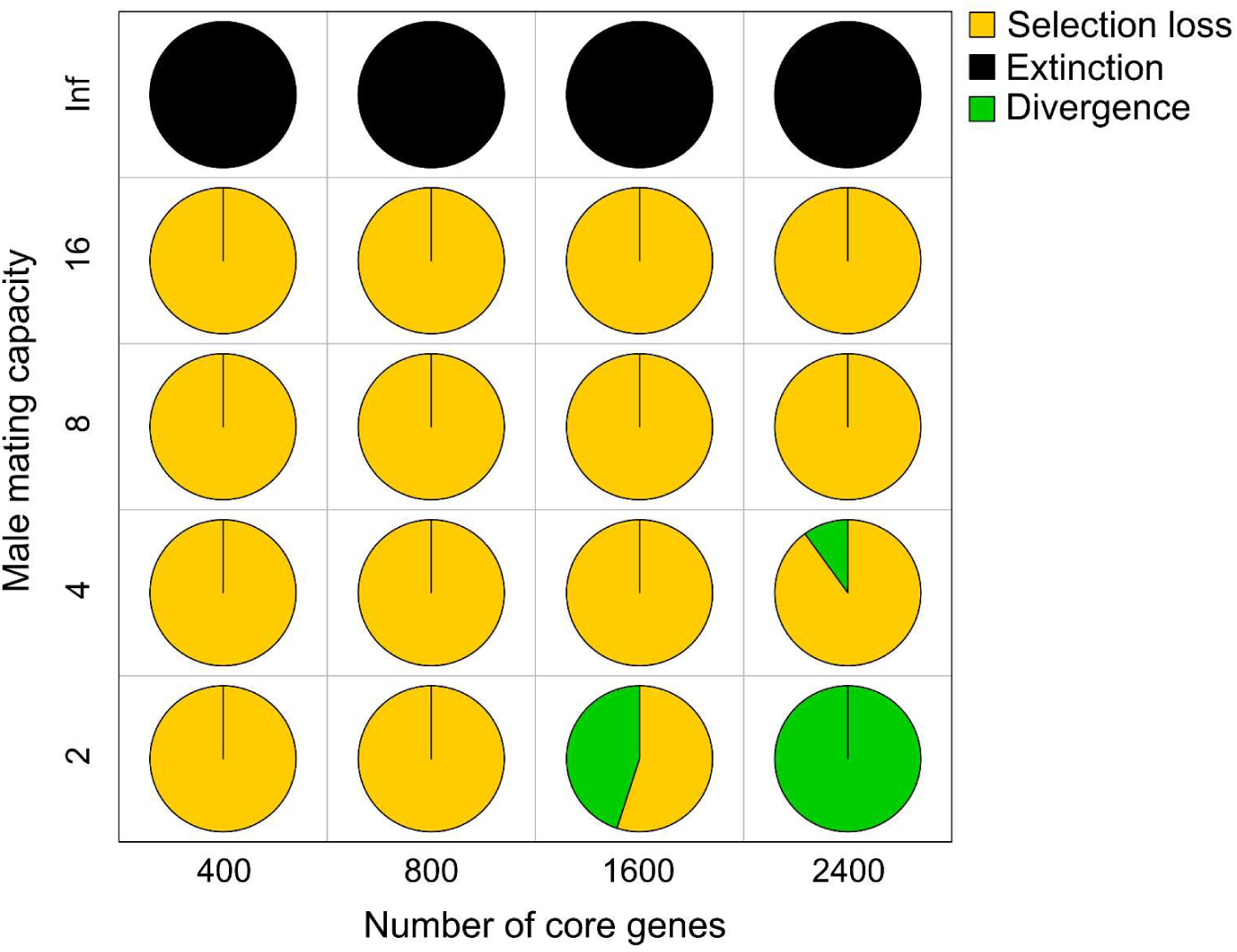
Simulation results under the additive genetic model with large values of *N* (the number of loci; horizontal axis) and varied values of *M* (the male mating capacity; vertical axis). Each pie chart shows frequencies of the following evolutionary outcomes of runaway across 20 simulation replicates under the same parameter conditions: cyclic evolution due to the loss of the selection gradient (yellow), ephemeral sympatric divergence (green), and evolutionary suicide (black). Other parameters were set to their default values (Table 1). Under the additive genetic model with large values of *N*, evolutionary dynamics were mostly cyclic due to the loss of the selection gradient, but these cyclic evolutionary dynamics sometimes led to an ephemeral sympatric divergence of the male display and female mate preference when *N* was very large and *M* was small (Figure S3).

**Figure S3.**
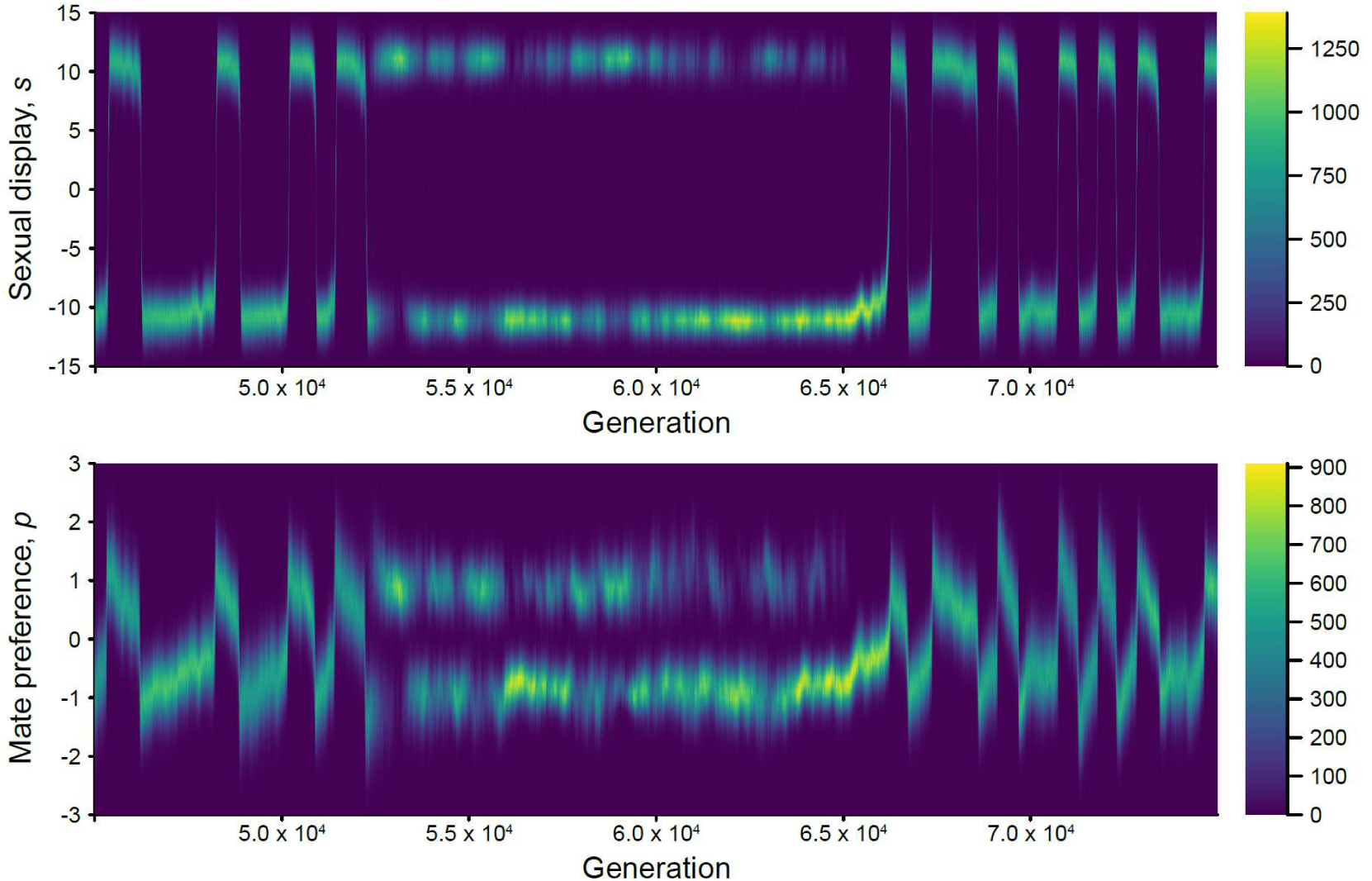
An example of an ephemeral sympatric divergence of the male display under an additive genetic model with 2000 loci. The phenotypic distributions of the male sexual display (*s*) and female mate preference (*p*) are shown for each generation (horizontal axis), with color intensity indicating the number of individuals. In this example, rapid cyclic evolutionary dynamics shifted into temporal coexistence of two extreme phenotypes during the period from generations 52,500–65,000. Similar dynamics have been observed in a previous model of sympatric speciation driven by Fisherian runaway (Higashi *et al*. 1999), in which runaway toward opposite extreme phenotypes can form two species, although they are unlikely to coexist stably (van Doorn *et al*. 2004). Parameters: *N* = 2000; *N* = 0; *N* = 0; *M* = 2; *θ* = 0.01; *K* = 5000. Other parameters were set to their default values (Table 1).

## Notes

### Competing Interest Statement

The authors have declared no competing interest.

